# Direct interaction between RSV polymerase L and active Rab11a mediates viral ribonucleoprotein transport to assembly sites

**DOI:** 10.1101/2025.10.28.685001

**Authors:** Claire Giry, Ana Joaquina Jimenez, Julie de Oliveira, Fatemeh Okhravijouybari, Mélissa Bessonne, Anne Huard de Verneuil, Patrick England, Priscila Sutto-Ortiz, Didier Chevret, Amina Tahir, Françoise Debart, Etienne Decroly, Julien Sourimant, Marie-Anne Rameix-Welti

**Affiliations:** M3P, Institut Pastetur, UMR 1173 (2I), Université Paris-Saclay, Université de Versailles Saint-Quentin, Inserm, Université Paris Cité, Paris, France; Unité de Virologie et Immunologie Moléculaires (VIM) UMR0892, INRAE-UVSQ-Université Paris-Saclay, Jouy-en-Josas, France; Plateforme de Biophysique Moléculaire, Institut Pasteur, Université Paris Cité, CNRS UMR3528, Paris, France; Aix Marseille Université, CNRS, AFMB UMR 7257, Marseille, France; IBMM, University of Montpellier, CNRS, ENSCM, Montpellier, France; Centre National de Référence Virus des Infections Respiratoires (CNR VIR), Institut Pasteur, Université Paris Cité, Paris, France

## Abstract

Respiratory syncytial virus (RSV) is an enveloped, negative-sense, single-stranded RNA virus whose ribonucleoproteins (vRNPs) must be transported from cytoplasmic viral factories to the plasma membrane for efficient virion assembly. Viral vRNPs comprise genomic RNA encapsidated by nucleoprotein N and associated with the polymerase complex (L, P, and M2-1). It was previously demonstrated that newly synthesized vRNPs are transported along microtubules by hijacking Rab11a, a small GTPase involved in the regulation of recycling endosomes. In our previous study, we showed an interaction between Rab11a and vRNPs in infected cells by immunoprecipitation assays, nevertheless the molecular mechanisms underlying Rab11a viral hijacking remained unknown.

Here, we provide the first comprehensive characterization of the interaction between RSV vRNPs and Rab11a using immunoprecipitation, immunofluorescence colocalization, GST pull-down assays, and biolayer interferometry. We demonstrate that the viral polymerase L is the sole vRNPs component responsible for Rab11a recognition: immunoprecipitation of L specifically co-precipitates HA-tagged Rab11a, whereas other vRNPs proteins show no interaction. *In vitro* binding studies confirm that L interacts directly and specifically with the active, GTP-bound form of Rab11a with sub-micromolar affinity. Domain mapping using truncated constructs reveals that this interaction requires the C-terminal methyltransferase and CTD domains of L (residues 1756-2165) and depends on Rab11a’s Switch I region, known to mediate interactions with cellular Rab11a partners. Mutagenesis further highlights leucine 1860 in the L polymerase as critical for Rab11a binding. Competitive inhibition of Rab11a-L interaction using the minimal Rab11a-binding domain reveals its involvement in vRNPs’ transport as it significantly impairs vRNP dynamics during infection. Together, these findings establish RSV polymerase L as the key mediator of Rab11a engagement, define the molecular interface of their interaction, and reveal a potentially conserved viral strategy for genome transport. Targeting the L-Rab11a interaction could therefore be a promising strategy for the development of RSV-specific or broad-spectrum antiviral therapies.

**Author Summary:** Respiratory syncytial virus is the leading cause of severe lower respiratory infection in children worldwide and is increasingly recognized as a major respiratory pathogen in the elderly and immunocompromised. Although vaccines have recently become available, treatment remains largely supportive in the absence of virus-specific antivirals, highlighting the urgent need for new therapeutic strategies.

In the infected cell, the viral genome is encapsidated and associated to the viral polymerase complex, to form viral ribonucleoproteins (vRNP). The vRNPs are produced in cytoplasmic viral factories and transported to the plasma membrane for assembly of new viral particles. Previous work has shown that RSV exploits Rab11a, a host GTPase that regulates recycling endosome trafficking, to mediate this transport. However, exactly how the virus connects to this transport system awaits to be precisely defined.

In this study, we identify the viral polymerase L as the key RSV protein that directly binds the active form of Rab11a. We mapped the domains of L and Rab11a that interact and showed that this connection is essential for efficient vRNPs trafficking.

These findings reveal a critical step in the RSV life cycle and suggest that disrupting the L-Rab11a interface could be a novel target for broad-spectrum antiviral development.

## Introduction

Human respiratory syncytial virus (RSV) is a highly contagious seasonal respiratory virus and the leading cause of acute lower respiratory tract infections (LRTIs) in young children [1,2]. Globally, RSV is estimated to cause about 33 million LRTIs each year, resulting in approximately 3.6 million hospitalizations and nearly 100,000 deaths among children under five [2]. RSV infects nearly all children by the age of two and is responsible for the majority of bronchiolitis and pneumonia cases during early childhood. During epidemic peaks, the associated symptomatology leads to the rapid saturation of pediatric intensive care units. RSV also poses a significant burden to the elderly and immunocompromised, resulting in substantial morbidity and mortality [3,4]. While effective preventive tools became available in 2023 - including maternal vaccines and long-acting monoclonal antibodies [5] for infants and vaccines for elderly [6,7] - treatment remains largely supportive in the absence of virus-specific antivirals, underscoring the continued need for new therapeutic targets.

RSV belongs to the order *Mononegavirales*, family Pneumoviridae, and genus Orthopneumovirus [8]. It is an enveloped virus with a non-segmented, negative-sense, single-stranded RNA genome of 15.2 kb encoding ten genes for eleven proteins. The genome is tightly encapsidated by the nucleoprotein N and associated with the polymerase complex - comprising the large polymerase L, phosphoprotein P, and transcription antiterminator M2-1 - forming viral ribonucleoproteins (vRNPs) that drive transcription and replication [9–11]. The L protein is a multidomain enzyme organized into distinct conserved regions: RNA-dependent RNA polymerase (RdRp), Polyribonucleotidyl transferase (PRNTase or capping enzyme), connecting domain, methyltransferase (MTase), and C-terminal domain (CTD), which together orchestrate viral RNA synthesis and processing [12,13]. The L protein remains tightly associated with P, which ensures L stability and contributes to its function [14].

Following membrane fusion and vRNP release into the cytoplasm, transcription and replication occur within membrane-less viral factories formed through liquid-liquid phase separation [15,16]. However, these factories are usually located far from the cell surface assembly sites, posing a critical transport challenge. Newly synthesized vRNPs are large, multi-megadalton complexes that cannot diffuse efficiently through the crowded cytoplasm, and they must travel substantial distances inside the cell to reach sites of virion formation [17]. This transport step thus represents a vulnerable point in the viral life cycle, as defective vRNPs trafficking would severely compromise viral spread.

To overcome this transport challenge, RSV hijacks cellular vesicular trafficking pathways by exploiting Rab11a-mediated recycling endosome transport. Rab11a belongs to the Rab11 subfamily of small GTPases, which in humans comprises three homologous isoforms: Rab11a, Rab11b (ubiquitously expressed), and Rab25/Rab11c (restricted to epithelial tissues). Rab11a functions as a molecular switch, cycling between active GTP-bound and inactive GDP-bound conformations [18]. Rab11a localizes primarily to the trans-Golgi network, post-Golgi vesicles, and apical recycling endosomes, where it regulates cargo sorting and polarized transport to the apical plasma membrane. In its active state, Rab11a recruits effector proteins (Rab11-FIPs) that engage motor proteins to drive vesicular transport along microtubules toward the plasma membrane [19]. Live-cell imaging has revealed that RSV vRNPs undergo rapid, directed movements along microtubules and colocalize with Rab11a. Under-expression of Rab11a abolished these directed vRNP movements, while co-immunoprecipitation experiments confirmed a physical interaction between Rab11a and vRNPs in infected cells [20]. Other viruses, as the influenza A virus, similarly exploit Rab11a for their own vRNPs trafficking [21,22]. Despite this established functional relationship, the molecular basis of RSV vRNP-Rab11a interaction remained completely unknown.

Here, we provide the first molecular characterization of the RSV-Rab11a interaction using a comprehensive combination of biochemical, biophysical, and cell biology approaches. Through analysis of individual vRNP components, we identify the large polymerase L as the sole viral component responsible for Rab11a recognition. Using purified recombinant proteins and biolayer interferometry, we demonstrate that L binds directly to Rab11a with submicromolar affinity, specifically recognizing the active GTP-bound conformation. Domain mapping reveals that the MTase and CTD regions of L are both necessary and sufficient for Rab11a binding, while mutagenesis of Rab11a shows that the Switch I region is essential for L recognition. We further identify leucine 1860 within the MTase domain as a potential residue contributing to this interaction. Functional validation using competitive inhibition demonstrates that disrupting the L-Rab11a interface significantly impairs vRNPs transport in infected cells, confirming the biological relevance of this molecular interaction.

## Results

### The interaction between vRNPs and Rab11a is mediated by the RSV Polymerase L

Previous studies have demonstrated that viral ribonucleoproteins (vRNPs) associate with Rab11a during transport. This interaction was confirmed by co-immunoprecipitation experiments in RSV infected cells [20]. To identify the viral protein that mediates vRNPs-Rab11a binding, we performed co-IP experiments using HEK-293T cells transiently co-expressing HA-Rab11a and one of the vRNPs proteins (N, P, L or M2-1) fused to GFP or BFP. We also co-expressed the matrix protein M - known to associate with vRNPs [23]. Prior to lysis, cells were treated with DSP (dithiobis(succinimidyl propionate)), a cross-linker that stabilizes protein-protein interactions. Cell lysates were incubated with agarose beads covalently linked to an anti-GFP nanobody (VHH) and bound proteins were revealed by western blotting (Figs 1A and 1B). Only the viral polymerase L enabled co-precipitation of HA-Rab11a when P, N, M2-1, and M failed to co-precipitate Rab11a. Although GFP-L was co-expressed with P to ensure its stability, Rab11a specifically co-immunoprecipitated with the L-P complex but not with P alone, indicating that L mediates the interaction with Rab11a. These results demonstrate that the viral polymerase L interacts with Rab11a and suggest that the association between RSV vRNPs and Rab11a is mediated by L.

**Fig 1.**
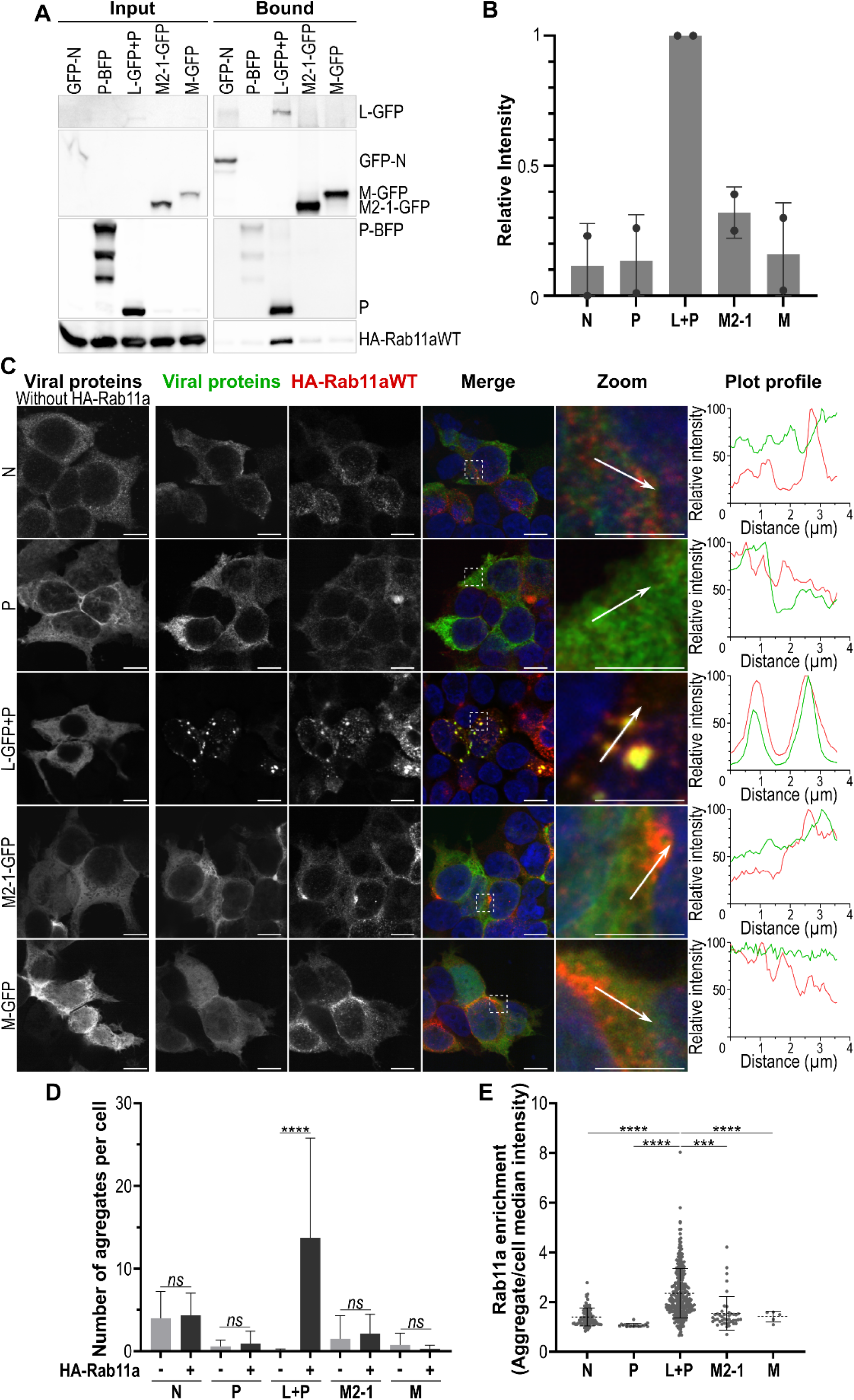
Viral polymerase L is the sole viral protein that interacts with Rab11a. HEK 293T cells were transfected with wild-type HA-Rab11a together with individual viral proteins (N, P, L, M2-1, or M) for 24h. **(A, B)** Co-immunoprecipitation analysis. Cells were treated with DSP cross-linking agent for 30 min, lysed and GFP-Trap beads were used to immunoprecipitate viral proteins as described in methods section. **(A)** Analysis of input lysates and precipitated fractions by Western blot using anti-GFP, anti-P, and anti-HA antibodies is shown. Representative of two independent experiments. **(B)** HA-Rab11a band intensities were quantified using ImageLab software and normalized to HA-Rab11a co-precipitated with L-GFP (ratio = 1). Bar graph shows mean ± SD from two independent experiments. **(C, D, E)** Immunofluorescence colocalization analysis. The indicated viral protein was expressed alone (left column) or together with HA-rab11a. Cells were fixed, immunolabelled and imaged using confocal microscopy to reveal viral proteins (green), HA-Rab11a (red) and nuclei (blue, Hoechst). **(C)** Images are representative of 3 independent experiments. Scale bars: 10 μm (main), 5 μm (zoom). Arrows indicate line scan vectors for intensity profiles. Fluorescence intensity is expressed as percentage of maximum signal detected for each condition. **(D)** Quantification of viral aggregate number per cell in absence or presence of HA-Rab11a co-expression. Bar graph shows mean ± SD from two independent experiments (n = 20 cells). *\*\*\*\* p* < 0.0001 by t-test with Welch’s correction. **(E)** HA-Rab11a enrichment in viral aggregates, calculated as ratio of HA-Rab11a intensity within aggregates to median cellular intensity. Dot plot shows mean ± SD, each point representing one aggregate. *\*\*\* p* < 0.0001, *\*\*\*\* p* < 0.0001 by Brown-Forsythe and Welch ANOVA test followed by Dunnett’s multiple comparisons test. Data from n = 20 cells, two independent experiments.

We further explored this interaction by analyzing the relative locations of Rab11a and the viral proteins using immunofluorescence microscopy. HEK-293T cells were transiently co-transfected with one expression plasmid encoding either N, P, L, M2-1 or M, alongside either an empty plasmid or a HA-Rab11a expression plasmid (Fig 1C). In the absence of Rab11a overexpression, all viral proteins as detected by specific antibody staining (N, P) or GFP fluorescence (L-GFP, M2-1-GFP, M-GFP) displayed diffuse cytoplasmic distribution. By contrast, upon Rab11a overexpression, the polymerase L underwent significant relocation, forming distinct cytoplasmic aggregates that colocalized with HA-Rab11a. While this aggregation represents an overexpression artifact not observed in infected cells, it provides a reliable readout for protein-protein interactions in subsequent experiments. The other viral proteins (N, P, M2-1-GFP and M-GFP) retained their diffuse cytoplasmic localization, regardless of the levels of Rab11a expression. Intensity profile analysis along defined cellular segments confirmed specific colocalization between L-GFP and HA-Rab11a, with overlapping fluorescence peaks not observed for other viral proteins. Quantitative analysis of aggregate formation (Fig 1D) revealed a significant increase in L-GFP-positive aggregates upon Rab11a overexpression compared to control conditions, while other viral proteins showed no significant changes in aggregation patterns. To assess the specificity of this interaction, we calculated Rab11a enrichment within viral protein aggregates (Fig 1E) as the ratio of HA-Rab11a fluorescence intensity within aggregates to the median cellular HA-Rab11a intensity. This analysis demonstrated specific enrichment of HA-Rab11a within L-GFP aggregates compared to the few aggregates formed by other viral proteins, confirming that L-GFP aggregates correspond to sites of L-Rab11a concentration.

Taken together, these results indicate that the viral polymerase L interacts with Rab11a, suggesting that it is likely the viral protein responsible for mediating the association between RSV vRNPs and this cellular trafficking factor.

### The polymerase L preferably interacts with the GTP-bound form of Rab11a

Rab11a is a small GTPase that cycles between two conformational states: an active, membrane-associated GTP-bound form, and an inactive, cytosolic GDP-bound form. To test whether the RSV polymerase L interacts preferentially with one of these states, we used two well-characterized Rab11a mutants: a constitutively active mutant Q70L (“CA”), locked in the GTP-bound state, and a dominant-negative mutant S25N (“DN”), locked in the GDP-bound state and unable to bind GTP [24]. Plasmids encoding HA-tagged Rab11a WT, CA, or DN were co-transfected with L-GFP in the presence of P. Co-immunoprecipitation assays were performed under cross-linking conditions, as described above (Fig 2A). Analysis revealed that L-GFP efficiently co-precipitated HA-Rab11a WT and HA-Rab11a CA, whereas HA-Rab11a DN was not detected in bound fraction. Quantification showed that Rab11a CA recovery was approximately twice that of Rab11a WT (Fig 2B) suggesting that the co-precipitated Rab11a WT represents the fraction of Rab11a in the active conformation. These data demonstrate that L specifically interacts with the active, GTP-bound form of Rab11a.

**Fig 2.**
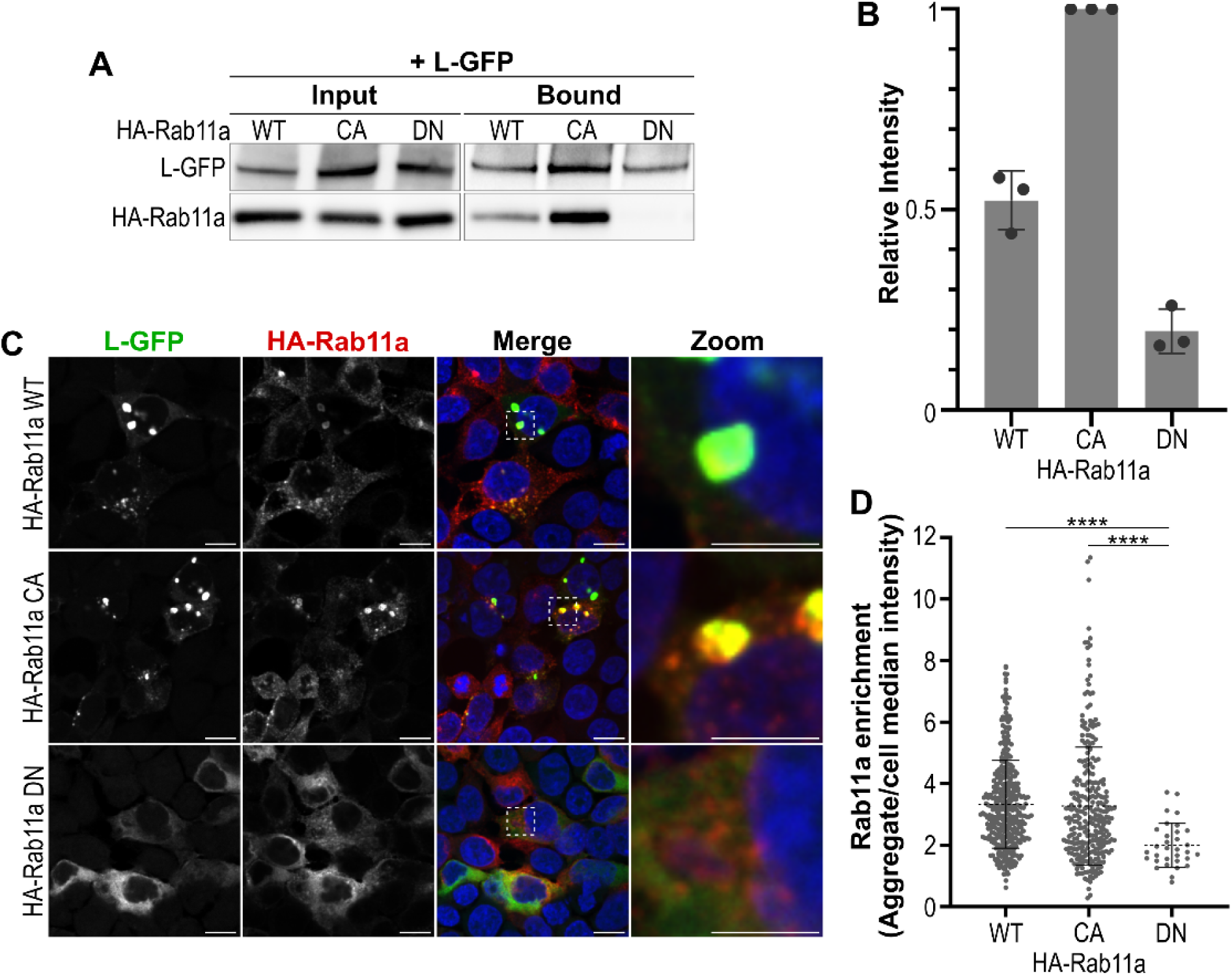
Polymerase L interact preferentially to the GTP-bound Rab11a. HEK 293T cells were transfected with wild-type (WT), constitutively active (CA, Q70L) or dominant negative (DN, S25N) HA-Rab11a together L-GFP and P for 24h. **(A, B)** Co-immunoprecipitation analysis. Cells were treated with DSP cross-linking agent for 30 min, lysed and GFP-Trap beads were used to immunoprecipitate L-GFP. **(A)** Analysis of input lysates and precipitated fractions by Western blot using anti-GFP, and anti-HA antibodies is shown. Representative of three independent experiments. **(B)** HA-Rab11a band intensities were quantified using ImageLab software and normalized to HA-Rab11a CA co-precipitated with L-GFP (ratio = 1). Bar graph shows mean ± SD from three independent experiments. **(C, D)** Immunofluorescence colocalization analysis. Cells were fixed, immunolabelled and imaged using confocal microscopy to reveal L-GFP (green), HA-Rab11a (red) and nuclei (blue, Hoechst). Scale bars: 10 μm (main), 5 μm (zoom). **(C)** Images are representative of three independent experiments. Quantification of viral aggregate par cell is shown in S1 Fig **(D)** HA-Rab11a enrichment in L aggregates, calculated as ratio of HA-Rab11a intensity within aggregates to median cellular intensity. Dot plot shows mean ± SD, each point representing one aggregate. *\*\*\*\* P* < 0.0001 by Brown-Forsythe and Welch ANOVA test followed by Dunnett’s multiple comparisons test. Data from n = 30 cells, three independent experiments.

We next assessed the subcellular distribution of Rab11a mutants relative to L using immunofluorescence microscopy. As observed previously, L-GFP expressed alone was diffusely distributed throughout the cytoplasm. When co-expressed with HA-Rab11a WT or HA-Rab11a CA, however, Rab11a and L relocalized into cytoplasmic aggregates (Fig 2C). In contrast, in the presence of HA-Rab11a DN, both proteins remained diffusely distributed without colocalization. To quantify aggregate formation, we measured the percentage of the total cell area occupied by L-GFP aggregates in each condition (S1 Fig). Cells co-expressing HA-Rab11a WT or CA displayed comparable fractions of their cytoplasmic area occupied by L-GFP aggregates, which was significantly larger than in cells expressing Rab11a DN. Consistently, Rab11a enrichment analyses (Fig 2D) demonstrated that HA-Rab11a WT and CA were specifically accumulated within L-GFP aggregates, whereas HA-Rab11a DN showed no enrichment.

Taken together, these results demonstrate that RSV polymerase L interacts preferentially with the GTP-bound, active form of Rab11a, consistent with the role of active Rab11a in transport of vRNPs.

### The polymerase L directly interacts with GTP-bound Rab11a

We next investigated whether the L-Rab11a interaction requires additional viral or cellular components or occurs through direct binding. To address this question, we performed GST pull-down assays using recombinant proteins. L-P^CTD^ complexes (P C-terminal Domain (CTD) is co-expressed to stabilize L) were expressed using the baculovirus system, while GST-tagged Rab11a WT, CA, and DN variants were produced in *E. coli*. Purified proteins were incubated together in the presence of Glutathione Sepharose 4B beads to capture GST-fusion proteins. Pull-down analysis revealed direct interaction between L-P^CTD^ and Rab11a CA or Rab11a WT in the presence of GTP. By contrast, Rab11a DN and Rab11a WT in the absence of GTP showed no detectable binding to L-P^CTD^ (Fig 3A). These results demonstrate that the L-Rab11a interaction is direct and requires Rab11a to be in its GTP-bound conformation.

**Fig 3.**
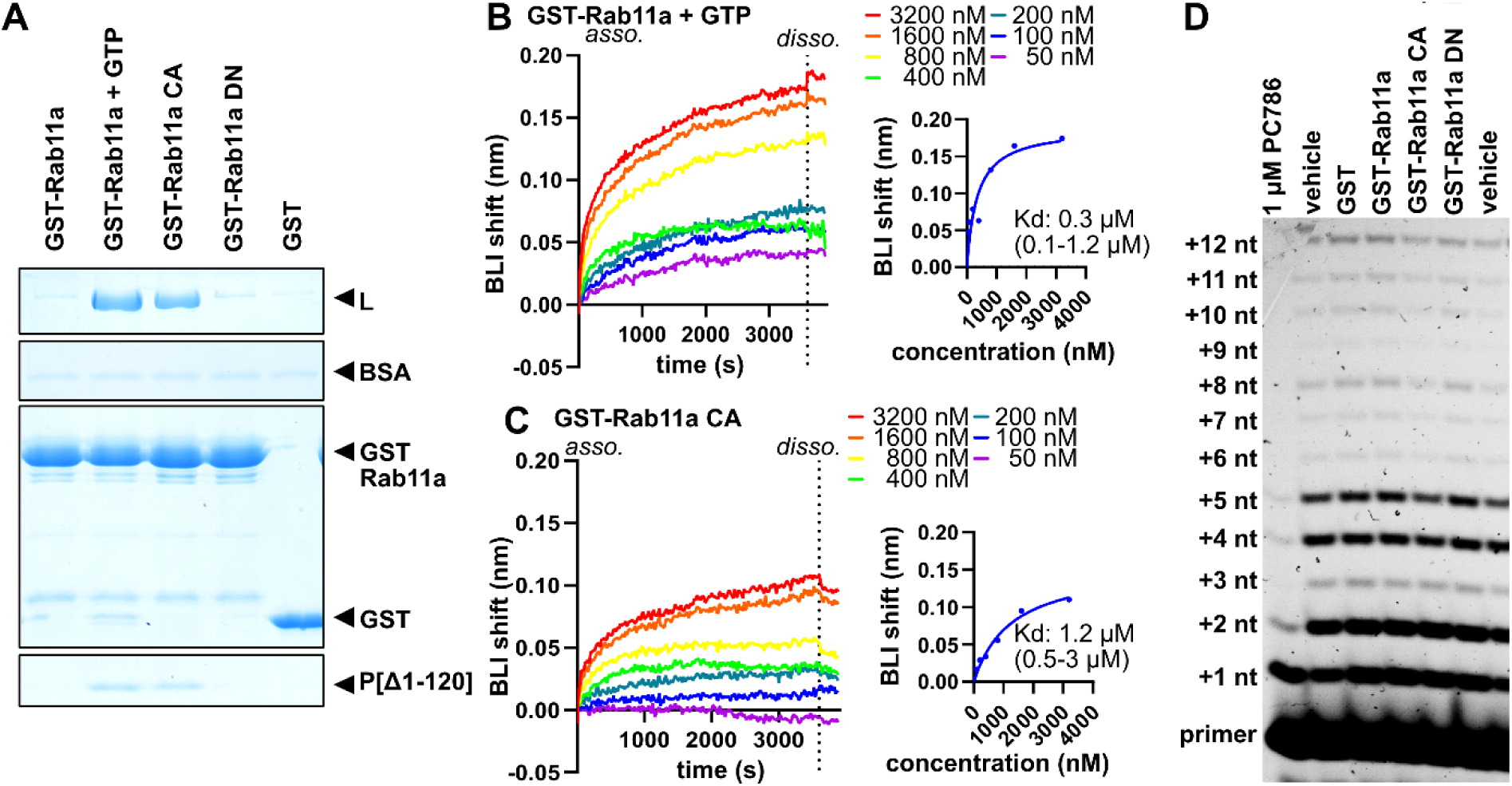
The polymerase L directly interacts with GTP-bound Rab11a. **(A)** Affinity coprecipitation with Glutathione Sepharose beads. Purified RSV L-Twin-Strep-tag + P[Δ1-120] was incubated with purified GST-Rab11a WT, GST-Rab11a(Q70L) CA, GST-Rab11a(S25N) DN and GST. After extensive washes, protein bounds were analyzed through Coomassie blue after SDS–polyacrylamide gel electrophoresis. Representative picture of five independent experiments. **(B,C)** Real-time biolayer interferometry profiles showing association and dissociation of GST-Rab11a (B) and GST-Rab11a CA (C) with RSV L-Twin-Strep-tag + P-his loaded on Ni-NTA sensors. **(D)** 3’ primer-extension assay. RSV L-Twin-Strep-tag + P-his was incubated with a fluorescently-labeled RNA primer annealed to a 16nt RNA template. The RdRp activity leads to accumulation of fluorescent RNA primers extended with 1-12 nucleotides. The RdRp activity was assayed in presence of GST-Rab11a WT, CA or DN. The activity in presence of PBS (vehicle) or GST was included as negative controls for inhibition, while an inhibitor (PC786, 1µM) of the RSV L RdRp activity was included as a positive control for inhibition. Fluorescent primers extended by the RSV polymerase were separated by Urea-PAGE and their fluorescence detected by a fluorescence imaging system. Representative gel from 3 independent experiments; quantification shown in S3B Fig.

To further characterize this interaction, we employed biolayer interferometry (BLI). Purified L-P^-his^ complexes were immobilized on Ni-NTA biosensors via the P^-his^ C-terminal His₆ tag and exposed to Rab11a variants. Initial experiments using single concentrations confirmed specific association of Rab11a WT in the presence of GTP and the constitutively active mutant, while no interaction was detected in the absence of GTP or with the DN mutant (S2 Fig). To determine the binding affinity, we exposed immobilized L-P complexes to increasing concentrations of either Rab11a (50-3200 nM) in the presence of GTP, or Rab11a CA (Figs 3B and 3C). Association kinetics were slow with equilibrium reached for all concentrations, enabling steady-state analysis. Fitting of equilibrium responses revealed specific interaction with an apparent KD in the sub-micromolar range for GST-Rab11a in presence of 1mM GTP (0.3 μM). Of note, minimal dissociation was observed during the buffer wash phase, indicating particularly stable binding under these *in vitro* conditions. However, this unusually high stability likely reflects the absence of physiological cofactors.

### Rab11a does not impact RSV polymerase activity

Having established that Rab11a directly interacts with the viral polymerase, we hypothesized that Rab11a binding might modulate L function, potentially inactivating the polymerase during intracellular transport. To test this possibility, we employed three complementary assays measuring distinct polymerase activities in the presence or absence of Rab11a variants.

We first assessed the impact of Rab11a on viral transcription and replication using an RSV minireplicon system [25]. HEK-293T T7 cells were transiently transfected with plasmids expressing components of the viral polymerase complex (N, P, L, M2-1) along with a pseudoviral RNA construct containing the firefly luciferase reporter gene. In this setup, the level of luciferase activity reflects the ability of the viral proteins to replicate and transcribe the pseudoviral RNA. Co-transfection with Rab11a variants (WT, CA, or DN) enabled us to evaluate polymerase activity in the presence of different Rab11a conformational states. Luciferase activity measurements revealed no significant differences between control conditions and cells expressing either Rab11a WT, CA, or DN (S3A Fig). These results indicate that Rab11a does not substantially affect the transcription and replication functions of the viral polymerase complex in this cellular context.

To directly evaluate the RNA-dependent RNA polymerase (RdRp) activity of L, we performed *in vitro* primer extension assays. In this system, purified L-P^-his^ complexes were provided with a synthetic RNA template annealed to a short fluorescently-labeled RNA primer, which is extended from the 3’ end by the polymerase using the template as a guide. Transcription products were visualized by denaturing gel electrophoresis. Gel analysis revealed distinct bands corresponding to synthesized RNA products with similar sizes and intensities across all conditions tested: GST control and GST-Rab11a fusion proteins (WT, CA, DN) (Fig 3D). Quantification of relative RNA transcript accumulation, expressed as percentage relative to the vehicle-treated control condition (set to 100%), confirmed that Rab11a variants had no detectable effect on L RdRp activity *in vitro* (S3B Fig).

Finally, we assessed the methyltransferase (MTase) activity embedded in the RSV L-P complex using a radioactive filter-binding assay. The RSV L protein contains MTase activity within its C-terminal domain, which catalyzes methylation of viral mRNA caps (both 2′-O and N7 methylation), essential for transcript stability and translation efficiency. This assay monitors the transfer of methyl groups from a radioactive donor, tritiated S-adenosyl methionine ([³H]-SAM), to a synthetic RNA substrate containing a cap structure analog (GpppRSV9) by purified L-P^-his^ complexes. Following the methylation reaction, RNA products are captured on DEAE filter mats while free donor molecules are removed by washing. The incorporated radioactivity directly reflects L MTase activity. Rab11a was used in excess at 0.5, 1.6, or 4.7 μM compared to the RSV L-P complex at 50 nM. This assay showed no significant differences in RSV polymerase MTase activity in the presence of purified Rab11a WT, CA, or DN compared to the GST control (S3C Fig). The methylation efficiency remained consistent across all tested conditions, indicating that Rab11a binding does not modulate the cap-modifying function of L.

Neither transcription, replication, RNA synthesis, nor cap methylation activities were significantly affected by the presence of Rab11a in any of its conformational states. Collectively, these results strongly suggest that Rab11a interaction does not result in inactivation of RSV polymerase enzymatic functions.

### The Switch I region of Rab11a is required for interaction with L

Having established that L specifically interacts with the GTP-bound form of Rab11a, we sought to identify the molecular determinants of this interaction. The Switch I region of small GTPases is a critical interaction surface for effector proteins and has been implicated in viral protein binding. Previous studies demonstrated that the Switch I motif of Rab11a, specifically the conserved IGV sequence (residues 44-46), is a docking site for influenza A virus polymerase PB2 [26]. As both the influenza A and RSV polymerases interact with Rab11a, we hypothesized that RSV polymerase L might utilize a similar binding interface on Rab11a. To test this hypothesis, we generated a Rab11a mutant in which the conserved IGV sequence was replaced with three alanines, creating the Rab11a_AAA_ mutant (Fig 4A). This triple alanine substitution is expected to disrupt the native structure of the Switch I region, while likely preserving the overall protein fold, allowing us to specifically assess the contribution of this motif to L binding.

**Fig 4.**
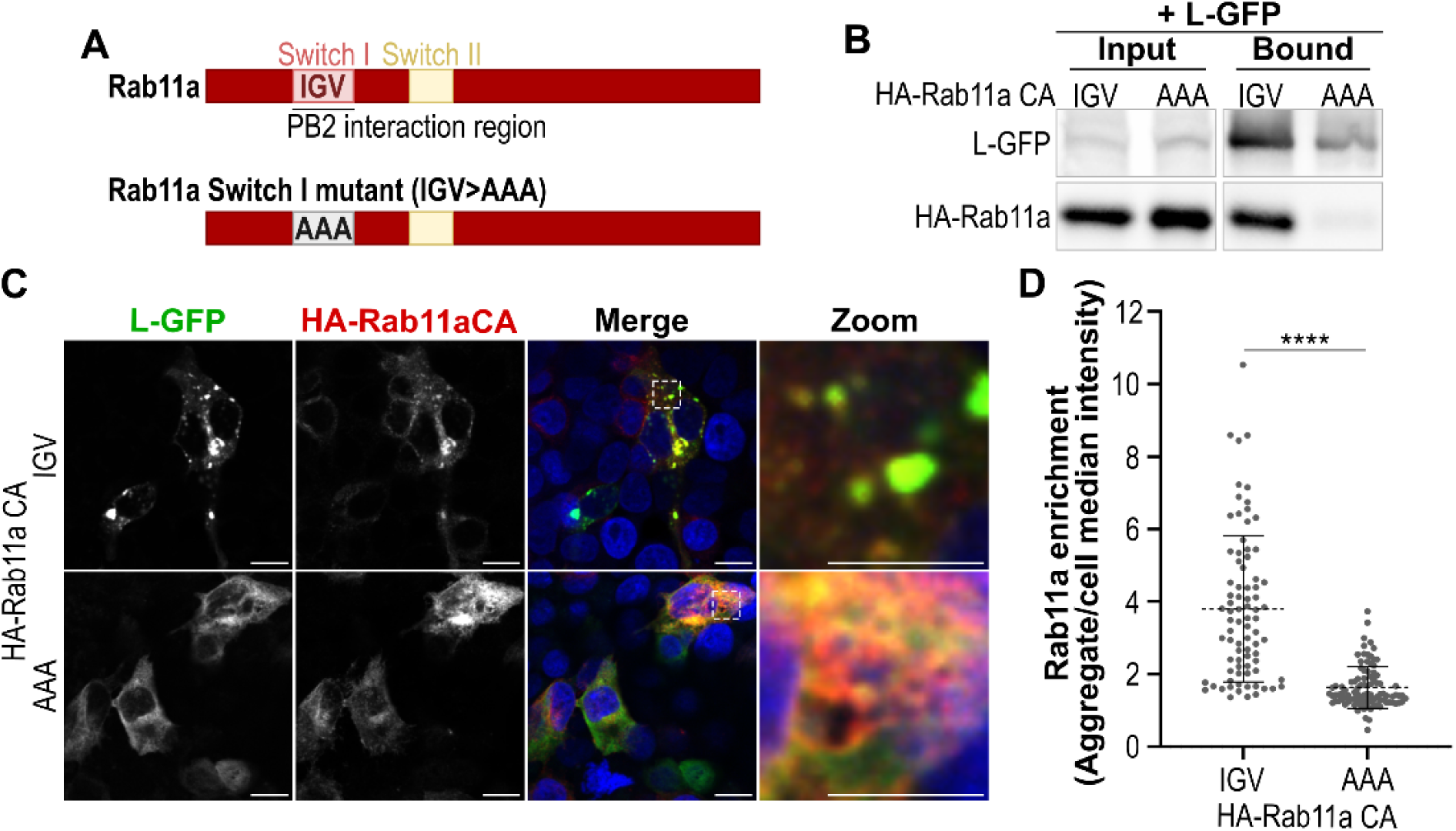
The Switch I region of Rab11a is essential for interaction with RSV L polymerase. **(A)** Schematic representation of Rab11a Switch I region mutation. Amino acids 44-46 were substituted from IGV to AAA. **(B, C, D)** HEK 293T cells were transfected with mutated (AAA) or unmutated (IGV) HA-Rab11a(Q70L) CA together with L-GFP and P for 24h. **(B)** Co-immunoprecipitation analysis. Cells were treated with DSP cross-linking agent for 30 min, lysed and GFP-Trap beads were used to immunoprecipitate L-GFP. Analysis of input lysates and precipitated fractions by Western blot using anti-GFP and anti-HA antibodies is shown. Representative of three independent experiments. Quantification of western blot signal is shown in S4A Fig. **(C, D)** Immunofluorescence colocalization analysis. Cells were fixed, immunolabelled and imaged using confocal microscopy to reveal L-GFP (green), HA-Rab11a (red) and nuclei (blue, Hoechst). **(C)** Images are representative of three independent experiments. Scale bars: 10 μm (main), 5 μm (zoom). Quantification of viral aggregate par cell is shown in S4B Fig. **(D)** HA-Rab11a enrichment in L aggregates, calculated as ratio of HA-Rab11a intensity within aggregates to median cellular intensity. Dot plot shows mean ± SD, each point representing one aggregate. *\*\*\*\* P < 0.0001* by unpaired t-test with Welch’s correction. Data from n = 20 cells, two independent experiments.

Co-immunoprecipitation assays revealed a complete loss of interaction between L-GFP and HA-Rab11aCA_AAA_ under cross-linking conditions (Figs 4B and S4A). While the HA-Rab11a CA signal in western blot analysis of the bound fraction was clear, only a very faint signal corresponding to background level was observed for HA-Rab11aCA_AAA_, despite equivalent input protein levels. This result demonstrates that the IGV motif within the Switch I region is essential for the L-Rab11a interaction, even when Rab11a is locked in the active, GTP-bound conformation.

To further validate these findings, we examined the subcellular distribution of the HA-Rab11aCA_AAA_ mutant relative to L using immunofluorescence microscopy. Consistent with our co-immunoprecipitation results, HA-Rab11aCA_AAA_ failed to induce L-GFP aggregation as L-GFP remained diffusely distributed throughout the cytoplasm upon HA-Rab11aCA_AAA_ overexpression (Fig 4C). Quantification of L-GFP aggregate formation revealed that cells expressing Rab11aCA_AAA_ displayed significantly reduced aggregate area compared to those expressing Rab11a CA (S4B Fig). Furthermore, Rab11a enrichment analysis showed no significant accumulation of HA-Rab11aCA_AAA_ within the few L-GFP aggregates that formed, confirming the loss of specific interaction (Fig 4D).

These results demonstrate that the Switch I region of Rab11a, specifically the conserved IGV motif, is absolutely required for interaction with RSV polymerase L.

### The two C-terminal domains of L are sufficient and essential for interaction with Rab11a

To further dissect the interaction from the viral side, we mapped the regions of L required for Rab11a binding. The RSV polymerase L is organized into five functional domains: the RNA-dependent RNA polymerase (RdRp), polyribonucleotidyl transferase (PRNTase), connecting domain (CD), methyltransferase (MTase), and C-terminal domain (CTD) (Fig 5A). We generated four truncated constructs to systematically test domain requirements: L_1-1460_-GFP, containing the RdRp and PRNTase domains, L_1-1755_-GFP, containing RdRp, PRNTase, and CD, GFP-L_1461-2165_, containing CD, MTase, and CTD, GFP-L_1756-2165_, containing MTase and CTD domains.

**Fig 5.**
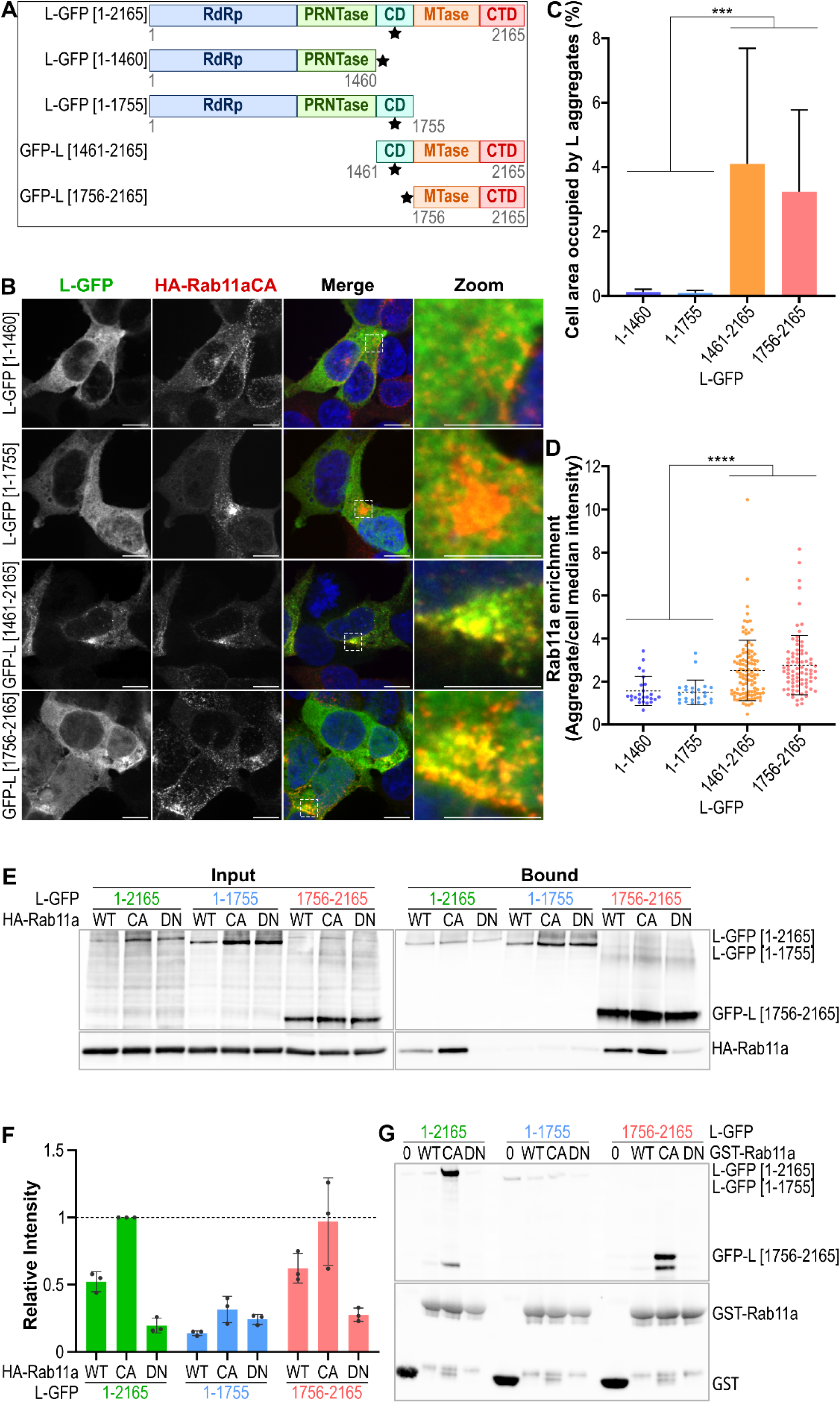
The C-terminal domains of RSV L are essential and sufficient for Rab11a interaction. **(A)** Schematic representation of L truncation constructs generated by deletion from full-length L (1-2165). Stars indicate GFP tag position. The tag is in the hinge 2 region of the connecting domain (CD) [15] or at the C-terminal end of L_1-1460_-GFP [27,28] **(B,C,D)** Immunofluorescence co-localization analysis. HEK 293T cells were transfected with HA-Rab11a CA together with P and L-GFP truncation constructs for 24h. Cells were fixed, immunolabelled and imaged using confocal microscopy to reveal L-GFP truncation constructs (green), HA-Rab11a (red) and nuclei (blue, Hoechst). **(B)** Images are representative of three independent experiments. Scale bars: 10 μm (main), 5 μm (zoom). **(C)** Quantification of cellular area occupied by L aggregates. Bar graph shows mean ± SD from two independent experiments (n = 20 cells). *\*\*\*\*P* < 0.0001 by Brown-Forsythe and Welch ANOVA test followed by Dunnett’s multiple comparisons test. **(D)** HA-Rab11a CA enrichment in L aggregates, calculated as ratio of HA-Rab11a intensity within aggregates to median cellular intensity. Dot plot shows mean ± SD, each point representing one aggregate. *\*\*\*\* P* < 0.0001 by Brown-Forsythe and Welch ANOVA test followed by Dunnett’s multiple comparisons test. Data from n = 20 cells, two experiments. **(E, F)** Co-immunoprecipitation analysis. HEK 293T cells transiently expressing HA-Rab11a WT, HA-Rab11a(Q70L) CA, HA-Rab11a(S25N) DN, together with P and L-GFP constructs for 24h, were treated with DSP for 30 min, lysed and incubated with GFP-Trap beads. **(E)** Analysis of input lysates and precipitated fractions by Western blot using anti-GFP, and anti-HA antibodies is shown. Representative of three independent experiments. **(F)** Quantification of co-immunoprecipitation. HA-Rab11a band intensities were quantified using ImageLab software and normalized to HA-Rab11a WT co-precipitated with full-length L-GFP (ratio = 1). Bar graph shows mean ± SD from three independent experiments. **(G)** GST pull-down assay. Sepharose beads coupled with GST or GST-Rab11a WT, CA or DN were incubated with lysates from HEK 293T cells transiently expressing L-GFP truncation constructs. Analysis of the precipitated fractions by Western blot using anti-GFP and anti-GST antibodies is shown. Representative of two independent experiments. Quantification is presented on S5A Fig.

We first screened these constructs for colocalization with Rab11a CA by immunofluorescence microscopy (Figs 5B-D). HA-Rab11aCA colocalized with the C-terminal fragments GFP-L_1461-2165_ and GFP-L_1756-2165_, but not with the N-terminal fragments L_1-1460_-GFP or L_1-1755_-GFP. To confirm these observations, we performed co-immunoprecipitation assays targeting GFP in cross-linked lysates of cells transiently expressing the truncated L proteins fused to GFP, together with P and HA-Rab11a variants (Figs 5E and 5F). Consistent with our immunofluorescence results and the behavior of full-length L (L1-2165), the C-terminal fragment GFP-L_1756-2165_ specifically co-precipitated HA-Rab11a WT and CA but not HA-Rab11a DN. In contrast, the N-terminal fragment L_1-1755_-GFP failed to co-precipitate HA-Rab11a in any conformational state, confirming that these domains are dispensable for Rab11a interaction.

To further validate these findings, we performed reciprocal GST pull-down assays using purified GST-Rab11a variants immobilized on glutathione beads incubated with lysates from cells expressing GFP-tagged L constructs (Figs 5G and S5A). Both full-length L-GFP and C-terminal fragment GFP-L_1756-2165_ were efficiently pulled down by GST-Rab11a CA but not by GST control or GST-Rab11a DN. No precipitation was observed with GST-Rab11a WT, possibly due to the absence of GTP in this experiment. Conversely, N-terminal fragments showed no detectable binding to any GST-Rab11a variant, confirming the specificity of the interaction for the C-terminal region of L. The same experiment performed for full-length L-GFP without P co-expression yielded identical results (S5B Fig), further confirming that P is dispensable for the L-Rab11a interaction.

These mapping studies demonstrate that the MTase and CTD domains of L (residues 1756-2165) are both necessary and sufficient for binding to Rab11a. The N-terminal domains (RdRp, PRNTase, and CD) spanning residues 1-1755 are dispensable for this interaction. Thus, the L-Rab11a interaction is mediated through the C-terminal portion of the polymerase.

### Residue L1860 of polymerase L is essential for Rab11a interaction

To further define the Rab11a-binding site within the L C-terminal region, we sought to identify specific residues that might be critical for this interaction. Given that both RSV L and the cellular effector Rab11FIP2 interact with the Switch I region of Rab11a, we hypothesized that these proteins might share similarities in their Rab11a-binding interfaces. Alignment of the Rab11a-binding domain of Rab11FIP2 with the minimal L fragment capable of Rab11a interaction (L_1756-2165_) revealed a region of sequence similarity located immediately downstream of the conserved GxGxG motif, a nucleotide-binding site essential for MTase activity conserved among *Mononegavirales* polymerases [28] (S6A Fig). We introduced a point mutation to replace leucine 1860 with alanine (L_L1860A_-GFP). This leucine is located at a central position within the Rab11a binding domain of Rab11FIP2 and is highly conserved. To ensure that the L1860A mutation did not induce a complete misfolding of the L, we first assessed the catalytic activity of the L_L1860A_ mutant using the minigenome assay. The L_L1860A_ mutant retained approximately 25% of wild-type polymerase activity in minigenome reporter assays (S6B Fig), indicating that the MTase domain is still enzymatically active.

The L_L1860A_ mutant showed a complete loss of Rab11a interaction in co-immunoprecipitation assays (Figs 6A and S6C). While wild-type L-GFP efficiently co-precipitated both HA-Rab11a WT and CA under cross-linking conditions, L_L1860A_-GFP failed to recover detectable levels of Rab11a in any conformational state, despite equivalent expression levels. Immunofluorescence analysis confirmed these biochemical observations (Fig 6B). In contrast to wild-type L-GFP, which formed distinct cytoplasmic aggregates upon co-expression with HA-Rab11a CA, the L_L1860A_ mutant remained diffusely distributed throughout the cytoplasm with no evidence of colocalization or aggregate formation with Rab11a (S6D and S6E Figs). These results demonstrate that leucine 1860 is essential for the L-Rab11a interaction and support the conclusion that the MTase domain of L, particularly the area surrounding L1860, is involved in specific binding of the viral polymerase.

**Fig 6.**
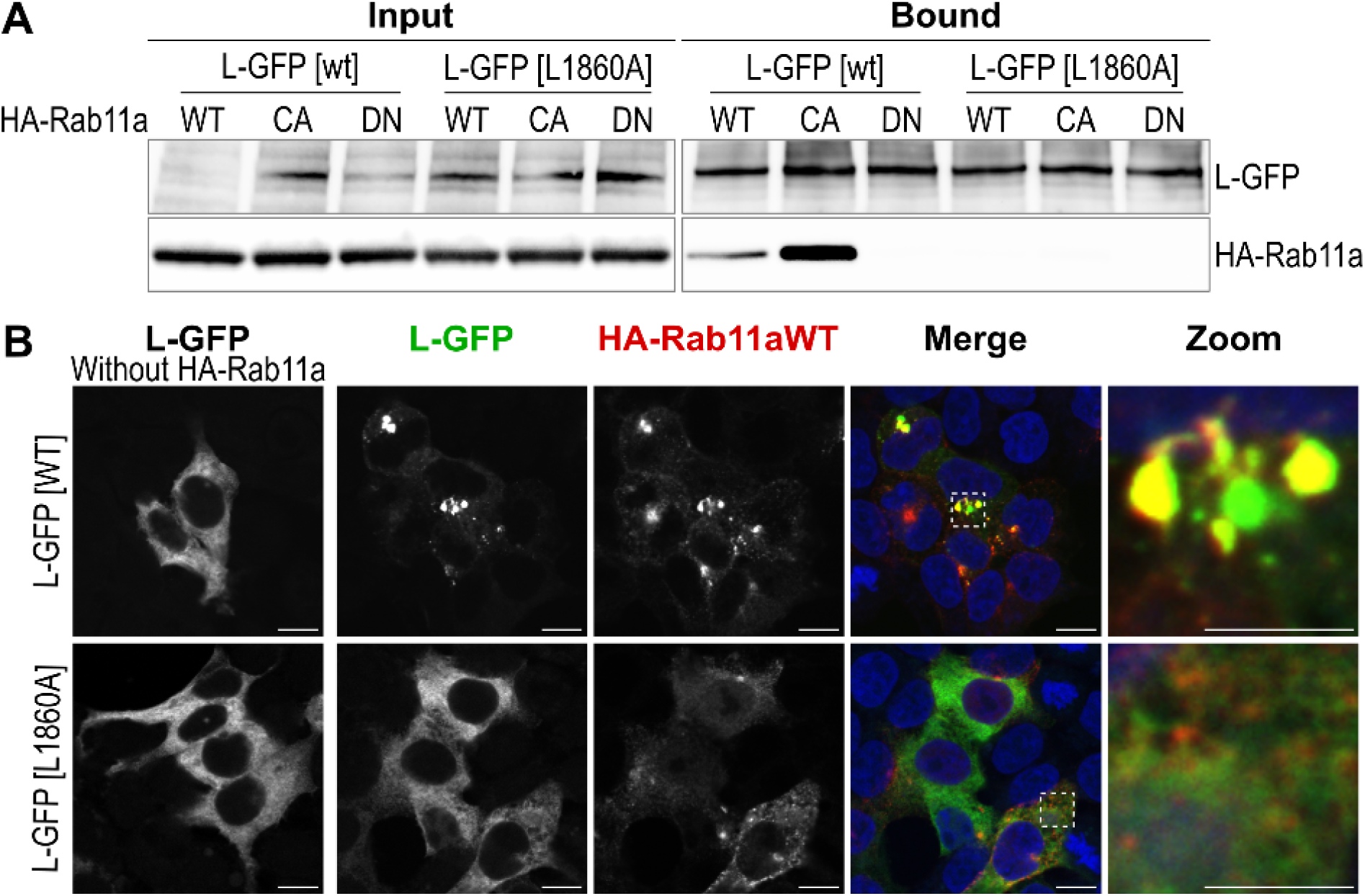
Leucine 1860 in polymerase L is important for Rab11a interaction. **(A)** Co-immunoprecipitation analysis. HEK 293T cells expressing HA-Rab11a WT, HA-Rab11a(Q70L) CA, HA-Rab11a(S25N) DN, together with P and L-GFP WT or L1860A mutant for 24h, were treated with DSP for 30 min 24 h post-transfection, lysed and incubated with GFP-Trap beads. Analysis of input lysates and precipitated fractions by Western blot using anti-GFP, and anti-HA antibodies is shown. Representative of two independent experiments. Quantification is presented on S6C Fig. **(B)** Immunofluorescence colocalization analysis. HEK 293T cells expressing HA-Rab11a CA together with P and L-GFP WT or L1860A mutant for 24h, were fixed, immunolabelled and imaged using confocal microscopy to reveal L-GFP WT or L1860A mutant (green), HA-Rab11a (red) and nuclei (blue, Hoechst). Scale bars: 10 μm (main), 5 μm (zoom). Images are representative of two independent experiments. Quantifications are presented on S6D and S6E Fig.

### The L-Rab11a interaction is required for efficient vRNPs transport in infected cells

Having identified and characterized the L-Rab11a interaction biochemically, we sought to determine its functional relevance in the context of RSV infection. To test whether this interaction is necessary for vRNPs transport, we designed a competitive inhibition approach using the minimal L fragment (L_1756-2165_) that binds to Rab11a as a dominant-negative inhibitor. We hypothesized that transient expression of mCherry-L_1756-2165_ would compete with the full-length viral L for binding to Rab11a, thereby disrupting (at least partially) vRNP-Rab11a association and possibly impairing vRNPs transport. To prevent potential toxicity from methyltransferase activity, we mutated the catalytic site within the MTase domain (R1820A and G1855S mutations [28]) to eliminate enzymatic function while preserving the Rab11a-binding capability. HEp-2 cells were transfected with either mCherry-L_1756-2165_ (competitive inhibitor) or mCherry alone (control), then infected with RSV-GFP-N to visualize vRNPs particles. At 18 hours post-infection, live-cell imaging was performed to track individual vRNPs movements over time as previously described in Consentino et al. [20] (See S7 and S8 Movies). Particle tracking analysis allowed quantitative assessment of multiple transport parameters including maximum speed, velocity and displacement.

Quantitative analysis of vRNPs tracking data revealed significant effects of L_1756-2165_ expression on vRNPs dynamics. While maximum speed (Fig 7A) showed only a modest, non-significant decrease, both track displacement (Fig 7B) and track velocity (Fig 7C) were significantly reduced in cells expressing the competitive inhibitor compared to mCherry controls. Visualization of centered track projections showed markedly reduced spatial coverage of vRNPs tracks in cells expressing mCherry-L_1756-2165_ compared to control cells, indicating less efficient long-range transport (Figs 7D and 7E). This spatial confinement suggests that vRNPs retain some motility but lose their ability to undergo sustained, directional transport.

**Fig 7.**
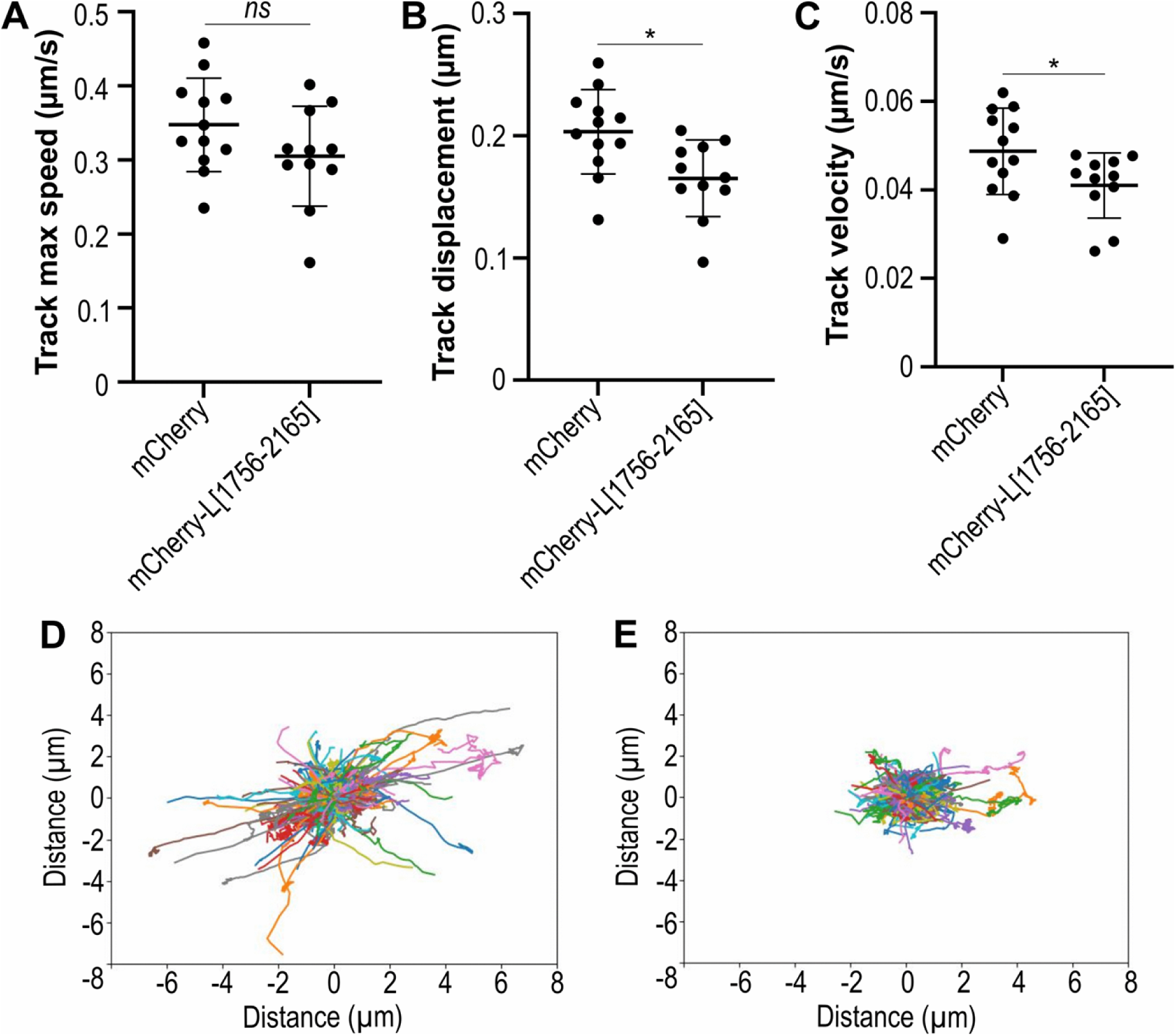
Long-range RNP transports depends on the L-Rab11a Interaction. HEp-2 cells were transfected with control mCherry or mCherry-L_1756-2165_ and infected with RSV-GFP-N. Live-cell imaging was performed at 18 h post-infection for vRNP tracking analysis (see method). **(A-C)** Quantification of RNP tracks parameters. Dot plots show mean ± SD and each data point represents the median values of track maximum speed **(A)**, track displacement **(B)**, and track velocity **(C)** from individual cells. *\*p* < 0.05 by unpaired t-test with Welch’s correction. Data from n = 12 cells across two independent experiments. **(D-E)** Representative centered projection of RNPs tracks in cells transfected with control mCherry **(D)** or mCherry-L_1756-2165_ **(E)** over 60 seconds. The corresponding time-lapse sequences are shown in S7 and S8 Movies.

These results provide functional evidence that the L-Rab11a interaction plays a critical role in vRNPs transport within infected cells. The moderate effects observed may reflect the competitive nature of our inhibition approach, where endogenous full-length L can still compete for Rab11a binding.

## Discussion

The transport of RSV vRNPs from viral factories to the plasma membrane occurs through rapid, directed movements that exploit host cellular machinery. These movements rely on the microtubule network and the small GTPase Rab11a [20]. Here, we present the first molecular characterization of the interaction between RSV ribonucleoproteins and the cellular trafficking factor Rab11a. We demonstrate that this interaction involves a direct association between the RSV polymerase L and Rab11a. Specifically, the two C-terminal domains of L (the MTase and the CTD, residues 1756–2165) bind to the Switch I region of Rab11a in its active, GTP-bound state.

Our analysis of individual viral proteins, conducted using immunoprecipitation and immunofluorescence techniques, demonstrates that polymerase L is the sole viral determinant responsible for recognizing Rab11a. In these experiments, we used a L protein fused with green fluorescent protein (GFP), with the GFP tag inserted just after the hinge 2 region of the connecting domain (position V1738) [29]. This fusion does not affect the polymerase’s functions, as evidenced by the ability to produce viruses that express L-GFP [30]. Confirmation of the interaction with purified, StrepTag-L protein shows that the GFP tag does not affect this interaction. Surprisingly, L-GFP aggregates when expressed in the presence of Rab11a overexpression. This aggregation is not observed in infected cells, possibly because L is recruited into VFs instead, and is thus not expected to have any role in viral replication. This phenotype is however a useful indicator of the interaction between L and Rab11a, enabling confirmation of IP data by an independent method. While P was co-expressed with L to ensure its stability, we can rule out the involvement of the co-expressed P or P^CTD^ in Rab11a binding. First, the known L-P interaction involves the core domain of L and the oligomerization/C-terminal domains of P [12,31], whereas the C-terminal fragment of L (L_1756-2165_) that mediates Rab11a binding does not interact with P. Moreover, GST pull-down experiments confirmed the L-Rab11a interaction in the absence of P.

The pull down and biolayer interferometry assays *in vitro* results demonstrate a direct interaction between L and Rab11a. At first glance, the binding affinity observed (0.3 μM) seems consistent with transient regulatory protein-protein interactions, which typically range from nanomolar to micromolar affinities [32,33]. Interestingly, we further observed that the association rate (kon) is actually very low, suggesting a potential conformational selection mechanism in which L or Rab11a may adopt a competent conformation before interacting. Dissociation is also very slow *in vitro*, suggesting a highly stable interaction in our assay possibly due to the locking of Rab11a in an active conformation. In a cellular context however, the interaction between L and Rab11a is expected to be further modulated by host regulatory factors, including guanine nucleotide exchange factors (GEFs) and GTPase-activating proteins (GAPs), which orchestrate the activation cycles of Rab11a [34,35].

We demonstrate that the Switch I region of Rab11a is essential for L recognition. This conserved region, along with Switch II, constitutes a key effector-binding interface of Rab11a, mediating interactions with diverse binding partners such as Rab11-FIP2 [36]. This raises the question of whether there is competition between L and other Rab11a effectors for binding to switch domain I of Rab11a, which could alter Rab11 function. The function of Rab11 is generally preserved in RSV infected cells [37], suggesting that either the pool of Rab11a involved is insufficient to have an impact or that there are redundancies. By contrast, Rab11 function is impaired in influenza A viruses infected cells, which could result from the interaction of the flu polymerase PB2 subunit with the switch domain of Rab11a.

As Rab11a is involved in vRNPs transport and has been shown to move with the vRNPs, it is tempting to speculate that the interaction between L and Rab11a mediates vRNPs movement. The expression of the minimal L domain (L_1756-2165_), which binds to Rab11a, significantly impaired vRNPs transport dynamics in infected cells, specifically affecting long, directed movements. However, vRNPs movements were not completely abolished, which is consistent with the possible involvement of Rab11 isoforms or alternative transport pathways. Furthermore, the L_1756-2165_ domain may only partially compete with the binding of the full-length L polymerase, as they may have a similar affinity for Rab11a. In any case, these data strongly suggest that this specific interaction is involved in the Rab11a-mediated transport of RSV vRNPs. One model is that newly synthesized vRNPs interact with Rab11a in its active GTP bound form (and therefore most likely associated with membranes) through the L domain and are transported with Rab11a vesicles along the microtubule network for long-range transport. Inactivation of Rab11 results in the release of vRNPs. In this model, preferential binding by L of the active form of Rab11a (bound to GTP) ensures that viral vRNPs only associate with transport-competent vesicles. However, it is unclear whether Rab11a only binds to L associated with vRNPs or whether it could also bind to free L, thereby interfering with its recruitment into VFs. In infected cells, or in cases of co-expression of N, P and L, L is clearly present in VFs, even under conditions of Rab11a overexpression. This suggests that the Rab11a-L interaction may be regulated. Our *in vitro* L-activity tests show that Rab11a has no effect on RNA synthesis activity, as determined by primer extension, or on methyltransferase activity in filter-binding assays. Moreover, minireplicon assay which provide a comprehensive readout of all RSV L activities including RNA synthesis, capping, and N7 methylation in a cellular context, demonstrate that L remains fully functional in the presence of Rab11a. In particular, interference by the Rab11a-L interaction, either on the capping or N7 methylation of luciferase mRNA, would result in decreased luciferase expression, which we did not observe. This is somewhat unexpected, particularly regarding methyltransferase activity, given that we have demonstrated Rab binding to the C-terminus of L (residues 1756-2165), including the MTase domain and the C-terminus domain. It is still possible that regulatory mechanisms exist to prevent Rab11a from recruiting a polymerase involved in viral RNA synthesis. One of these mechanisms could be the compartmentalization of RNA synthesis inside VFs. Several other key questions remain. For example, the precise timing of L-Rab11a engagement and the motor proteins (dynein and kinesin) that drive vRNP movement during the viral life cycle are unclear, as are the additional cellular factors that modulate this transport.

In recent years, Rab11a has emerged as a critical host factor for vRNPs trafficking across diverse viral families. Influenza A virus was the first virus shown to require Rab11a for vRNPs transport to assembly sites [21,22]. Rab11a has since been implicated in Ebola virus VP40 transport and VLP release, as well as in Sendai virus polymerase - mediated vRNPs recruitment to recycling endosomes [38,39]. Newly synthesized vRNPs from measles, mumps, parainfluenza viruses 1 and 3, Marburg virus, and New World hantaviruses have also been shown to associate with Rab11-positive endosomes and rely on Rab11 for transport to the plasma membrane [40–43]. The molecular mechanisms underlying virus-Rab11a interactions have been characterized for several viruses, revealing both conserved and virus-specific recognition strategies. In the case of influenza A virus, the export of vRNP depends on a direct protein-protein interaction between the C-terminal domain of the viral polymerase subunit PB2 and the Switch I region of GTP-bound Rab11a [21,26,44]. For Sendai virus, the viral polymerase complex (comprising L and its cofactor C) mediates vRNP recruitment to Rab11-positive recycling endosomes, although the precise binding interface and whether the interaction is direct or involves cellular adaptors remains to be determined [39]. Interestingly, human parainfluenza virus type 3 employs a distinct strategy, where the nucleocapsid protein N seems to mediate Rab11a recognition in its GTP-bound active form [45]. For measles, mumps, and hantavirus, while vRNP association with Rab11-positive compartments has been documented, the precise viral binding partners and molecular determinants remain unidentified.

The hijacking of the Rab11 pathway by numerous viruses, especially respiratory viruses, makes it an attractive target for broad-spectrum antivirals. A recent study showed that inhibiting Rab11 function with a peptide affected the replication of PIV3, RSV and influenza viruses [45]. However, given the importance of this pathway in cellular physiology, it may be preferable to block the interaction between the virus and Rab11 rather than the pathway itself. A better definition of the interaction domains is essential for this approach to be successful. The identification of leucine 1860 as a potentially important residue provides the first glimpse into the specific molecular contacts governing this interaction. However, it is unclear whether this residue is directly involved in interaction with Rab11A, or whether the mutation can locally alter the structure, resulting in a loss of binding. Indeed, L1860 is located in proximity to the GxGxG nucleotide-binding motif in the methyltransferase domain, and the MTase structure from HMPV (closely related to RSV) suggests that L1860 could be located underneath surface residues. This would support the hypothesis that the mutation may locally perturb the protein conformation. However, the conservation of partial enzymatic activity in L demonstrates that the MTase is still functional, ruling out a significant change in the conformation of this region and supporting the hypothesis that the surrounding region is involved in the interaction. Higher-resolution structural approaches such as cryo-EM or crystallography will be essential to define the binding interface at atomic resolution and to unlock rational design of targeted inhibitors. Interestingly, L1860 is highly conserved across the Mononegavirales order, particularly among viruses known to exploit the Rab11 pathway during infection. Sequence alignment shows that this leucine residue is found in the Ebola virus, Sendai virus, measles virus, parainfluenza virus 3, and Marburg virus (see S9 Fig). This conservation suggests that similar Rab11a-binding strategies may be widely used across this viral order. Further comparative studies are needed to determine whether polymerase C-terminal domains represent a universal feature.

In conclusion, this study establishes the molecular basis for RSV vRNPs trafficking through Rab11a-mediated transport. We demonstrate that the viral polymerase L directly recognizes active Rab11a through specific molecular determinants involving the MTase/CTD region of L and the Switch I region of Rab11a. This interaction is functionally required for efficient vRNPs transport during infection, representing a critical step in the viral life cycle. These findings allow our understanding of negative-strand RNA virus pathogenesis to progress significantly and provide a foundation for developing novel antiviral strategies targeting viral transport mechanisms.

## Materials and methods

### Cells

Human embryonic kidney 293T (HEK-293T) cells and HEK-293T T7 cells, stably expressing T7 RNA polymerase, were maintained in Dulbecco’s Modified Eagle Medium (DMEM), and human epidermoid larynx carcinoma HEp-2 cells (ATCC: CCL-23) were cultured in Eagle’s minimum essential medium (MEM). Both media were supplemented with 10% (v/v) heat-inactivated fetal bovine serum (FBS) and 1% (v/v) penicillin-streptomycin solution (5,000 U/mL). Cells were grown in an incubator at 37°C under 5% CO₂. Spodoptera frugiperda (SF9; ATCC, CRL-1711 Gibco™ cat# 11496015) were propagated in suspension using Sf-900 II serum-free medium (Gibco™ cat# 10902096) at 28°C without CO_2_.

### Virus

The recombinant respiratory syncytial virus RSV-GFP-N is derived from the human RSV Long strain sequence and was previously described [20]. It was amplified on HEp-2 cells at 37°C (3 passages) and titrated by plaque assay on HEp-2 [46].

### Plasmids

The pM/Luc subgenomic replicon containing the firefly luciferase (Luc) gene and expression plasmids pCI-N, pCI-GFP-N, pCI-P, pCI-P-BFP and pCI-M2-1-GFP have been previously described [15,47,48]. Expression plasmids pCI-L and pCI-M were constructed by cloning mammalian codon-optimized RSV L and M coding sequences into the pCI vector (GenBank U47119). Briefly, coding sequences were amplified by PCR using Phusion High-Fidelity DNA Polymerase (Thermo Fisher Scientific) with specific primers and cloned into pCI using MluI/XbaI restriction sites for L and MluI/NotI for M. To generate fluorescent fusion constructs, the GFP coding sequence was inserted into the Hinge 2 region of L as previously described [29] or at the C-terminus of M sequence, yielding pCI-L-GFP and pCI-M-GFP expression vectors, respectively. Truncated L constructs were generated from pCI-L-GFP by restriction digestion followed by Gibson assembly (New England Biolabs) as detailed in S10 Table. Briefly, pCI-L-GFP was digested with appropriate restriction enzymes. The GFP tag and deleted L coding sequences were PCR-amplified using Phusion High-Fidelity DNA Polymerase (Thermo Fisher Scientific) with primers designed to create overlapping ends. For constructs retaining the connector domain (pCI-L_1-1755_-GFP and pCI-GFP-L_1461-2165_), the eGFP tag remained in its original Hinge 2 position. For constructs lacking the connector domain (pCI-L_1-1460_-GFP and pCI-GFP-L_1756-2165_), the eGFP coding sequence was repositioned respectively at the C-terminus or N-terminus. PCR products were assembled with digested vectors using the Gibson Assembly Master Mix (New England Biolabs). The L1860A point mutation was introduced into pCI-L-GFP by site-directed mutagenesis using the In-Fusion HD Cloning Kit (Takara Bio). Mutagenic primers were designed to replace leucine 1860 with alanine (L1860A) and included 15 bp regions of homology flanking the mutation site. The entire plasmid was amplified using Phusion High-Fidelity DNA Polymerase (Thermo Fisher Scientific). The parental template plasmid was digested with DpnI. The linearized plasmid was recircularized using the In-Fusion HD Cloning Kit according to the manufacturer’s instructions. All constructs were verified by Sanger sequencing.

The pHA-Rab11a WT, pHA-Rab11a CA (Q70L), and pHA-Rab11a DN (S25N) expression plasmids were generated by replacing the N-terminal GFP tag with an HA epitope tag in peGFP-Rab11a constructs (kindly provided by Dr. Nathalie Sauvonnet). The parental peGFP-Rab11a plasmids (WT, CA, or DN) were digested with NheI/XhoI (WT) or NheI/SalI (CA and DN) to excise the eGFP coding sequence. Complementary oligonucleotides encoding the HA epitope tag (YPYDVPDYA) with compatible overhangs were synthesized, annealed, and ligated into the linearized vectors. The Rab11a Switch I mutant (I44A/G45A/V46A, designated Rab11a_AAA_) was generated from pHA-Rab11a CA by site-directed mutagenesis (In-Fusion HD Cloning Kit (Takara Bio)) to replace the conserved IGV sequence with three alanines as described above. All constructs were verified by Sanger sequencing. All primer sequences used in this study are available upon request.

For the purification of GST-tagged Rab11a proteins, the reference sequence for human Rab11a (uniprot identifier P62491) was codon-optimized for expression in *Escherichia coli* and a DNA fragment containing this sequence was synthesized (Twist Bioscience), alongside sequences variants encoding the CA (Constitutively Active) variant Q70L or DN (Dominant Negative) variant S25N. These sequences were inserted in pGEX 4T-3 expression plasmids (Invitrogen) multiple cloning sites (between EcoRI and XhoI restriction sites) downstream of a GST sequence, a thrombin-cleavage site sequence (LVPRGS), and short flexible linker sequence (PNSGGGGSGGGGS) using the In-Fusion HD Cloning Kit (Takara cat# 639650) to form a pGEX-Rab11a, pGEX-Rab11a CA and pGEX-Rab11a DN plasmids. All plasmids were amplified and then validated by Sanger sequencing.

For the purification of RSV P-L complexes, we modified a previously described pFastBacDual plasmid allowing the co-expression of both a codon-optimized RSV L protein with an HA-tag (YPYDVPDYASLGGP) at position 1740 under the control of the polyhedrin promoters, and a codon-optimized P protein with an N-terminal GST-tag under the control of the p10 promoters [31]. Using the In-Fusion HD Cloning Kit (Takara cat# 639650), the L protein sequence was extended at the C-terminus with a precision protease cleavage site (LLEVLFQGP) and a Twin-Strep-tag (SAWSHPQFEKGGGSGGGSGGSAWSHPQFEK), while the P protein N-terminal GST-tag was excised and its C-terminus was extended with a tobacco etch virus cleavage site (GSENLYFQ) and a 6xhis tag (HHHHHH), resulting in a pFBD RSV L-Twin-Strep-tag + P-his-tag plasmid. In parallel, we used this method to construct a pFBD RSV L-Twin-Strep-tag + P[Δ1-120] plasmid that allows the expression of a truncated P sequence that only contain the minimum P sequence that is required for L protein stabilization, as previously reported [49].

### Cell Transfection and Infection

Cells were seeded 24 hours prior to transfection to reach 90% confluence at the time of transfection in either 24-well plates (immunofluorescence), Ibidi μ-Slide 4-well chambers (live imaging), or 10 cm dishes (co-immunoprecipitation). Transfections were performed using Lipofectamine 2000 (Thermo Fisher Scientific) at a ratio of 2.5 μL per 1 μg of DNA, with DNA amounts adjusted to culture vessel size (0.5 μg for 24-well plates/μ-Slides; 10 μg for 10 cm dishes), in antibiotic-free growth medium, according to the manufacturer’s instructions. Cells were fixed or collected at 24 hours post-transfection (hpt) for subsequent analyses. For transfection-infection assays, cells were transfected as described above in MEM supplemented with 2% FBS without antibiotics. At 1 hpt, RSV viral stocks diluted in the same medium were added to the cells at multiplicity of infection (MOI) of 1. Cells were incubated at 37°C under 5% CO₂ and processed at 18 hours post-infection.

### Immunofluorescence and colocalization Analysis

At 24 hours post-transfection, cells were fixed with 4% paraformaldehyde in PBS for 15 minutes, permeabilized with PBS containing 1% BSA and 0.1% Triton X-100 for 10 minutes, and incubated for 1 hour at room temperature with primary antibodies diluted in PBS-BSA 1%: rabbit anti-RSV N (VIM Unit, INRAE, 1:40,000), rabbit anti-RSV P (VIM Unit, INRAE, 1:1,000) [50], or mouse anti-HA (BioLegend, clone 16B12, 1:800). After two washes with PBS, cells were incubated for 1 hour with Alexa Fluor 488- or 647-conjugated secondary antibodies against rabbit or mouse IgG (H+L) (Thermo Fisher Scientific, 1:1000), phalloidin-Atto-590 (Sigma-Aldrich, 1/800), and Hoechst 33342 (1 μg/mL). Coverslips were mounted with ProLong Diamond antifade reagent (Thermo Fisher Scientific). Representative cells were captured by confocal microscopy using an inverted spinning-disk confocal microscope (IXplore SpinSR, Evident, Tokyo, Japan) equipped with a 60× oil-immersion objective lens (Plan-Apochromat, Evident) and an ORCA-Fusion BT camera (C15440, Hamamatsu Photonics, Japan). Z-stacks were acquired and maximum intensity projections (1 μm) were generated using ImageJ (NIH). For colocalization analysis, viral protein-positive aggregates were automatically detected in co-transfected cells using threshold-based segmentation in ImageJ, allowing quantification of aggregate number per cell and surface area. Rab11a enrichment within aggregates was calculated as the ratio of mean Rab11a fluorescence intensity in viral protein-positive regions to the median Rab11a intensity in the entire cell. All image analyses were performed using custom ImageJ macros. Statistical analyses and data visualization were performed using Prism (GraphPad).

### Cross-linking and GFP Trap co-immunoprecipitation

Transfected cells were incubated in 300 mM DSP (dithiobis(succinimidyl propionate), Thermo Fisher Scientific) in PBS for 30 minutes at room temperature. The amine-crosslinking reaction was quenched with 20 mM Tris-HCl (pH 7.5) in PBS for 15 minutes at room temperature. Cells were then collected and lysed in 500 µL/10^7^ cells of buffer containing 25 mM Tris-HCl (pH 7.5), 200 mM NaCl, 1 mM EDTA, 0.5% NP40, 1 mM PMSF, and a cocktail of protease inhibitors (Thermo Scientific Pierce) on a rotary wheel at 4°C for 1 hour. Lysates were cleared by centrifugation (10,000 g, 10 minutes at 4°C) and incubated with GFP-Trap beads overnight on a rotary wheel at 4°C according to the manufacturer’s protocol. Beads were washed three times with lysis buffer before being resuspended in NuPAGE LDS sample buffer containing NuPAGE sample reducing agent (Thermo Fisher Scientific).

### Co-immunoprecipitation assay using GST-tagged proteins

GST-Rab11a conjugated beads were prepared before use : Glutathione Sepharose 4B beads (Cytiva) were incubated with purified GST-Rab11a fusion proteins (purification described below) for 1 hour at room temperature with constant rotation. Briefly, 5 mg of purified GST-fusion proteins were mixed with 1 mL of pre-equilibrated beads in binding buffer (50 mM Tris-HCl pH 7.5, 300 mM NaCl, protease inhibitor cocktail). Efficient protein loading onto beads was verified by SDS-PAGE analysis followed by Coomassie blue staining. Transfected cells were harvested and lysed in ice-cold lysis buffer containing 50 mM Tris-HCl pH 7.5, 300 mM NaCl, protease inhibitor cocktail, 0.1 mg/mL BSA, 1 mM DTT, 1 mM PMSF, and 0.6% NP-40. Cells were incubated in lysis buffer for 1 hour at 4°C with constant rotation to ensure complete lysis. Following centrifugation at 12,000 × g for 10 minutes at 4°C to remove cellular debris, clarified lysates were incubated with GST-Rab11a conjugated beads for 3 hours at 4°C under constant rotation. After binding, beads were washed three times with wash buffer (50 mM Tris-HCl pH 7.5, 300 mM NaCl, protease inhibitor cocktail) to remove non-specifically bound proteins. Bound protein complexes were eluted by resuspending washed beads in NuPAGE LDS sample buffer containing NuPAGE sample reducing agent (Thermo Fisher Scientific).

### Western Blot

Samples were denatured at 80°C for 10 minutes, separated by SDS-PAGE, and transferred to PVDF membranes using liquid transfer systems according to standard protocols. Membranes were blocked for 1 hour at room temperature in PBS containing 0.25% Tween-20 (PBS-T) and 5% non-fat dry milk, then incubated overnight at 4°C with primary antibodies diluted in PBS-T containing 1% BSA: rabbit anti-RSV P (VIM Unit, INRAE, 1:1,000), rat anti-HA (Roche, 1:800), mouse anti-GFP (Roche, 1:2,000), or HRP-conjugated mouse anti-GST (BioLegend, 1:1,000). After washing with PBS-T, membranes were incubated for 1 hour at room temperature with appropriate HRP-conjugated secondary antibodies for mouse, rabbit, or rat IgG (H+L) (1:20,000; Promega or Santa Cruz Biotechnology). Proteins were visualized using SuperSignal West Pico PLUS or Atto Ultimate chemiluminescence substrates (Thermo Fisher Scientific) on a ChemiDoc imaging system (Bio-Rad). Densitometric quantification was performed using Image Lab software (Bio-Rad).

### Protein Expression and Purification

The RSV L-P polymerase complex was expressed and purified using a protocol based on a previously described Sf9/baculovirus system [31] with the following modifications. Bacmids allowing for the expression of a yellow fluorescent protein and carrying either the RSV L-Twin-Strep-tag + P-his-tag sequences (shortened as L-P^-his^ in the main text) or the RSV L-Twin-Strep-tag + P[Δ1-120] sequences (shortened as L-P^CTD^ in the main text) were recovered after transformation of the corresponding pFBD plasmids in the DH10EMBacY bacteria (Geneva Biotech). The bac-to-bac baculovirus expression system (Invitrogen) was used to generate recombinant baculoviruses in SF9 cells by transfection with Cellfectin II (Gibco) reagent following the manufacturer’s instructions. Baculovirus stocks were amplified twice at MOI of 0.01 in SF9 cells to generate high titer baculovirus stock (1-3*10^8^ TCID50/ml). Titration was performed by 50% tissue culture infectious dose (TCID50) using yellow fluorescence to monitor infection.

For L-P protein production, SF9 cells in suspension were infected at a MOI of 1 for 72h. Infected cells were harvested by centrifugation at 300 g for 10 mins and lysed in a lysis buffer containing 50 mM Tris-HCl pH 8.0, 300 mM NaCl, 0.5% Igepal, 1 mM DTT, 2.5 U/ml Benzonase nuclease and an antiproteases and antiphosphatases cocktail (Pierce) for 1h on ice. Lysates were clarified by centrifugation 20 mins at 20,000xg at 4°C. The L-P complex was purified using a two-step chromatographic approach. Cell lysate was first applied to a Strep-Trap XT column (Cytiva) and the L-Twin-Strep-tag was eluted with biotin-containing buffer (50 mM Tris-HCl pH 8.0, 300 mM NaCl, 1 mM DTT, 50 mM Biotin). The eluate was then concentrated on Vivaspin 6 100kDa (Sartorius) and subjected to size exclusion chromatography on a Superose 6 increase 10/300 GL column (Cytiva) pre-equilibrated with 50 mM Tris-HCl pH 8.0, 300 mM NaCl, 1 mM DTT. Fractions containing the L and P proteins were collected and further concentrated on Vivaspin 6 100kDa (Sartorius) to ∼0.5-1 mg/ml, flash-frozen and stored at -80°C. Proteins were stored in 50 mM Tris-HCl pH 8.0, 300 mM NaCl, 1 mM DTT buffer at 0.6 mg/ml at -80°C.

Wild-type and mutant GST-Rab11a fusion proteins (GST-Rab11a, GST-Rab11a CA, and GST-Rab11a DN) along with GST control were expressed after transformation of corresponding pGEX 4T-3 plasmids in *Escherichia coli* BL21 cells. Bacterial cultures were selected by ampicillin (100 µg/ml) and grown at 37°C to an OD_₆₀₀_ of 0.6, then induced with 0.33 mM IPTG for 16 hours at 28°C.

Cells were harvested by centrifugation and resuspended in lysis buffer containing 50 mM HEPES pH 7.4, 500 mM NaCl, 10% glycerol, 0.1% octyl β-D-glucopyranoside, 1 mM DTT, 1 mg/mL lysozyme, protease inhibitor cocktail (Roche), and 2.5 U/mL Benzonase nuclease following a previously described protocol [26]. Cell debris after sonication were removed by centrifugation at 15,000 × g for 30 minutes at 4°C. The soluble fraction was applied to Glutathione Sepharose 4B FastFlow resin (Cytiva). After extensive washing, bound proteins were eluted with 50 mM reduced glutathione in 50 mM HEPES pH 7.4, 500 mM NaCl. Eluted fractions were further purified by size exclusion chromatography using a Superdex 200 increase 10/300 GL column pre-equilibrated with PBS. Peak fractions were pooled, and protein concentrations were determined using the Bradford assay with bovine serum albumin as standard. Final protein concentrations ranged from 2-5 mg/ml, and aliquots were flash-frozen and stored at -80°C. A quality control of the protein samples was conducted in accordance with the ARBRE-MOBIEU guidelines [51].

### GST Pull-Down Assay

Recombinant GST, GST-Rab11a wild-type, GST-Rab11a CA, and GST-Rab11a DN fusion proteins were normalized to equimolar concentrations (35 μM) in PBS. Glutathione Sepharose 4B beads (20 μl settled volume per reaction) were pre-equilibrated in binding buffer (50 mM Tris-HCl pH 8.0, 300 mM NaCl) and incubated with 4 µM of each GST fusion protein and 0,6 µM of RSV L-Twin-Strep-tag + P[Δ1-120] supplemented with 0.1 mg/ml BSA and 0.02% Tween 20 for 1 hour at room temperature with constant shaking at 1300 rpm. For nucleotide-dependent binding studies, GTP was included at a final concentration of 1 mM. Following binding, beads were washed twice with binding buffer containing 0.1 mg/ml BSA and 0.02% Tween 20, then twice with binding buffer alone. When applicable, GTP (1 mM) was maintained throughout all wash steps. Bound proteins were eluted in 2× SDS sample buffer and resolved by SDS-PAGE and visualized by Coomassie blue staining. Input controls consisting of individual GST constructs and L-P complex were included in all experiments.

### Biolayer interferometry

Protein-protein interactions and binding affinities between RSV L-Twin-Strep-tag + P- his complexes and GST-Rab11a variants were analyzed by biolayer interferometry (BLI) using an Octet RED 384 system at 25°C with orbital shaking at 1,000 rpm. Purified RSV L-P complexes and GST-Rab11a variants were buffer-exchanged into assay buffer containing 50 mM Tris-HCl pH 8.0, 300 mM NaCl, 0.02% Tween-20, 100 μg/mL or 1 mg/mL BSA and 1 mM GTP when specified, using PD-10 Desalting Columns (Cytiva). For Protein-protein interaction assays, RSV L-Twin-Strep-tag + P-his complexes (20 μg/mL) were loaded onto Ni²⁺-NTA biosensors via the C-terminal 6×His tag on the P protein for 900 seconds. Streptavidin-His6 coated sensors were used as controls to assess non-specific sensor binding. After baseline establishment in assay buffer (60 seconds), loaded biosensors were exposed to a 2 µM concentration of GST-Rab11a variants for 600s (association) and to assay buffer for 300s (dissociation step). As control for signal drift, loaded sensors were incubated in assay buffers in parallel. For binding affinity measurements, loaded sensors were incubated in two-fold serial dilutions of GST-Rab11a or GST-Rab11a CA fusion proteins ranging from 50 nM to 3.2 μM (two-fold increments). Association was monitored for 60 minutes until binding reached steady state. Then biosensors were incubated in assay buffer for 5 min (dissociation). BLI data were processed using Octet Data Analysis software. Since minimal dissociation was observed, equilibrium dissociation constants (KD) were determined by fitting steady-state binding responses to a one-site binding model using GraphPad Prism software, after double referencing of the BLI response by subtracting non-specific sensor binding and baseline drift signals.

### Minigenome assay

Cells in 24-well plates were transfected as described above with a plasmid mixture containing the following RSV minigenome system components per well : 200 ng pCI-N, 200 ng pCI-P, 100 ng pCI-L or variant constructs (pCI-L-GFP, pCI-L L_1860A_-GFP), 50 ng pM2-1, 200 ng pCI-GFP or pHA-Rab11a variants (WT, CA or DN), and 200 ng pGEM-M/Luc containing the RSV minigenome cassette with firefly luciferase reporter gene. To normalize for transfection efficiency, 10 ng of pβ-Gal (expressing β-galactosidase under CMV promoter control) was included in each transfection. Negative control transfections were performed by replacing pCI-L with empty pCI vector. All experimental conditions were performed in biological triplicate, with each independent experiment repeated three times. At 24 h post-transfection cells were lysed in luciferase lysis buffer (25mM Tris pH 7.9, 8mM MgCl_2_, 1mM DTT, 1% Triton X-100 and 15% glycerol) for 5 minutes at 37°C. Luciferase activity was measured using a Centro XS³ LB 960 luminometer (Berthold Technologies) and normalized on β-galactosidase activity.

### RNA synthesis

RNA primer (5′-[6-FAM]ACCA-3′) and RNA template (5’-CUUUGUUUUUUCUGGU-3’) were chemically synthesized on a solid support at 1 µmol scale using an ABI 394 automated synthesizer with commercial 2’-*O*-pivaloyloxymethyl 3’-*O*-phosphoramidite ribonucleosides and 6-FAM phosphoramidite (Chemgenes, USA). After assembly of RNA sequences, the solid support was directly subjected to 1M DBU in dry CH_3_CN for 3 minutes then 30% aqueous ammonia solution for 3h at 40 °C to recover the crude RNA material. After filtration, evaporation in the presence of isopropylamine, and coevaporation with water, the two short RNA substrates were purified by IEX-HPLC and their identity was confirmed by MALDI-TOF mass spectrometry. 191 nmoles of pure RNA template and 442 nmoles of pure RNA primer were obtained.

### *In vitro* polymerase assay

RNA polymerase activity was assessed using a primer extension assay based on a previously described assay, with a modification: radiolabeled nucleotides were omitted and instead the RNA primer was fluorescently labeled in order to visualize transcription products by fluorescence [27]. Each reaction (5 μL final volume) contained 0.2 μM recombinant RSV L-Twin-Strep-tag + P-his complex, 7 μM GST fusion proteins (GST control, GST-Rab11a WT, GST- Rab11a CA, or GST-Rab11a DN), 2 μM RNA template (3′-UGGUCUUUUUUGUUUC-5′), and 100 μM fluorescent RNA primer (5′-[6-FAM]ACCA-3′). Reactions were performed in buffer containing 50 mM Tris-HCl pH 8.0, 60 mM NaCl, 8 mM MgCl₂, 1 mM TCEP, 10 μM each NTP, and 10% glycerol. As a control, a reaction containing the RSV L inhibitor PC786 at 1 µM was included in a separate reaction [52]. Following incubation for 3 hours at 30°C, reactions were terminated by adding one volume (5 µl) of dionized formamide with 25 mM EDTA and denatured at 95°C for 3 minutes. RNA products were resolved on 20% polyacrylamide gels containing 7 M urea and visualized using a GelDoc Go imaging system (Bio-Rad).

### MTase activity assay

Methyltransferase activity was measured using a filter-binding assay performed according to the method described previously [53]. Reactions contained 50 nM of RSV L-P complex, GST fusion proteins at varying concentrations (0.5, 1.6, or 4.7 μM), 1.8 μM synthetic RNA substrate (GpppRSV_9_: 5′-GpppGGGACAAAA-3′), 0.17 μM unlabeled S-adenosyl methionine (SAM), and 0.8 μM [³H-methyl]-SAM (PerkinElmer) in reaction buffer (50 mM Tris-HCl pH 8.0). After 3 hours incubation at 30°C, reactions were quenched by a 20-fold dilution in cold water. Samples were transferred to DEAE filtermats (PerkinElmer) using a Filtermat Harvester (Packard Instruments) to separate methylated RNA products from radiolabelled SAM. The RNA-retaining mats were washed twice with 10 mM ammonium formate pH 8.0, twice with water and once with ethanol. They were then soaked with scintillation fluid (PerkinElmer), and ^3^H-methyl transfer to the RNA substrates was determined using a Wallac MicroBeta TriLux Liquid Scintillation Counter (PerkinElmer).

### Live Imaging and vRNPs Tracking

Time-lapse images were acquired for 1 min at 37 °C and 5% CO₂ using an inverted spinning-disk confocal microscope (IXplore SpinSR, Evident, Tokyo, Japan) equipped with a 60× oil-immersion objective lens (Plan-Apochromat, Evident) and an ORCA-Fusion BT camera (C15440, Hamamatsu Photonics, Japan). The system was fitted with a CSU-X1 confocal scanning unit and a SORA disk module (Yokogawa, Tokyo, Japan), coupled to a 3.1× relay lens. System control and image acquisition were performed using CellSens software (Molecular Devices, Sunnyvale, CA, USA). Live-cell imaging and particle tracking analyses were performed as previously described [20]. Briefly, Z-stacks were acquired at intervals of less than 200 milliseconds and processed using ImageJ for maximum projections and background subtraction. Particle tracking was conducted using Imaris software 10.2 (Bitplane Inc.) with the autoregressive motion algorithm. The data were further filtered and analyzed using a custom Python script, which is available at https://github.com/mawelti/RSV-RNP-TrackAnalysis. Statistical comparisons between medians were performed using Prism software (GraphPad Inc.).

## Acknowledgments

We thank Sandrine Rosario and the Molecular Biophysics platform of Institut Pasteur for quality control analysis of purified proteins. We are grateful to Dr. Nathalie Sauvonnet for Rab11a plasmids. This work was supported by the French National Research Agency (ANR) under the project RSVFact, grant number ANR-21-CE15-0030-01 (MARW), and ReSVIC, ANR-24-CE44-2004 (JS).

## Supporting information

**S1 Fig. Polymerase L interacts preferentially with GTP-bound Rab11a.** Quantification of cellular area occupied by L aggregates corresponding to Fig 2C. Bar graph shows mean ± SD from three independent experiments (n = 30 cells). *\*\*\*\*P* < 0.0001 by Brown-Forsythe and Welch ANOVA test followed by Dunnett’s multiple comparisons test.

**S2 Fig. The polymerase L directly interacts with GTP-bound Rab11a.** Association of GST-Rab11a(Q70L) CA and GST-Rab11a(S25N) DN (left graph) or Rab11a WT with or without GTP (right graph) at fixed 2 µM concentration with either immobilized RSV L-Twin-Strep-tag + P- his-tag complex or immobilized Streptavidin-His6 protein as a negative control, on Ni-NTA biosensors. Representative sensorgrams showing association (asso) and dissociation (disso) kinetics.

**S3 Fig. Rab11a interaction has no impact on RSV L activity. (A)** Polymerase activity assay in presence of Rab11a WT, CA or DN mutants. HEK 293T T7 cells were transfected with plasmids encoding N, P, L, M2-1, and M/Luc subgenomic minireplicon together with control GFP or HA- Rab11a WT, HA-Rab11a(Q70L) CA, HA-Rab11a(S25N) DN. Polymerase activity was measured by luciferase luminescence normalized to β-galactosidase activity. Data represent mean ± SD from two independent experiments performed in triplicate. **(B)** Densitometry analysis of the intensity of selected fluorescent RNA transcripts in the 3’ primer extension assay shown in Fig 3D. **(C)** *in vitro* methyltransferase (MTase) activity assay. The RSV L-Twin-Strep-tag + P-his-tag Methyltransferase activity was assayed in the presence of GST-Rab11a WT, CA or DN *in vitro* using an established protocol [53]. As a control, a known inhibitor of RSV methyltransferase activity (Sinefungin) was included.

**S4 Fig. The Switch I region of Rab11a is essential for interaction with RSV polymerase L. (A)** Quantification of co-immunoprecipitation shown in Fig 4B. HA-Rab11a band intensities were quantified using ImageLab software and normalized to unmutated (IGV) HA-Rab11aCA (ratio = 1). Bar graph shows mean ± SD from three independent experiments. **(B)** Quantification of cellular area occupied by L aggregates corresponding to Fig 4C. Bar graph shows mean ± SD from two independent experiments (n = 20 cells). *\*\*\*P* < 0.0005 by unpaired t-test with Welch’s correction.

**S5 Fig. The C-terminal domains of RSV polymerase L are necessary and sufficient for Rab11a interaction. (A)** Quantification of GST pull-down shown in Fig 5G. L-GFP truncation construct band intensities were quantified using ImageLab software and normalized to full-length L-GFP co-precipitated with GST-Rab11aCA (ratio = 1). Bar graph shows mean ± SD from two independent experiments. **(B)** GST pull-down assay. Sepharose beads coupled with GST, GST- Rab11a WT, GST-Rab11a(Q70L) CA or GST-Rab11a(S25N) DN were incubated with lysates from HEK 293T cells transiently expressing L-GFP with or without P. Analysis of the precipitated fractions by Western blot using anti-GFP, anti-P and anti-GST antibodies is shown. One experiment.

**S6 Fig. Leucine 1860 in polymerase L is important for Rab11a interaction. (A)** Sequence alignment of L_1756-2165_ with the Rab11a-binding domain (RBD) of Rab11-FIP2 using Clustal Omega. Leucine 1860 is framed in red. Symbols indicate conservation levels: (*) fully conserved residue; (:) conservation between amino acids with strongly similar properties; (.) conservation between amino acids with weakly similar properties. **(B)** Polymerase activity assay. HEK 293T T7 cells were transfected with plasmids encoding the N, P, L-GFP WT or L1860A mutant, M2-1, and the M/Luc subgenomic minireplicon. Polymerase activity was measured by luciferase luminescence normalized by the to β-galactosidase activity as described in methods. Bar graph shows mean ± SD from four independent experiments performed in triplicate. **(C)** Quantification of co-immunoprecipitation shown in Fig 6A. HA- Rab11a CA band intensities were quantified using ImageLab software and normalized to HA- Rab11a CA co-precipitated with WT L-GFP (ratio = 1). Bar graph shows mean ± SD from two independent experiments. **(D,E)** Quantifications of immunofluorescence colocalization analysis shown in Fig 6B. **(D)** Quantification of cellular area occupied by L aggregates. Bar graph shows mean ± SD from two independent experiments (n = 20 cells). *\*\*\*\*P* < 0.0001 by unpaired t-test with Welch’s correction. **(E)** HA-Rab11aCA enrichment in L aggregates, calculated as ratio of HA-Rab11a intensity within aggregates to median cellular intensity. Dot plot shows mean ± SD, each point representing one aggregate *\*\*\*\*P* < 0. 0001 by unpaired t-test with Welch’s correction. Data from n = 20 cells, two experiments.

**S7 Movie. RNP dynamics in control mCherry-transfected infected cells.**

Time-lapse microscopy of vRNPs in Hep-2 cells transfected with control mCherry and infected with RSV-GFP-N. Imaging was performed at 18-20 h post-infection using spinning disk confocal microscopy with 0.19 second intervals for 1 minute. Video was processed using ImageJ software (10 frames per second) after maximum z-projection. Representative video (related to Fig 7D) from two independent experiments. Scale bar: 5 μm.

**S8 Movie. RNP dynamics in mCherry-L_1756-2165_-transfected infected cells.**

Time-lapse microscopy of vRNPs in Hep-2 cells transfected with control mCherry-L_1756-2165_ and infected with RSV-GFP-N. Imaging was performed at 18-20 h post-infection using spinning disk confocal microscopy with 0.19 second intervals for 1 minute. Video was processed using ImageJ software (10 frames per second) after maximum z-projection. Representative video (related to Fig 7E) from two independent experiments. Scale bar: 5 μm.

**S9 Fig. Sequence alignment of MTase domains of viruses known to exploit Rab11 during infection using Clustal Omega.** Leucine 1860 is framed in red. Symbols indicate conservation levels: (*) fully conserved residue; (:) conservation between amino acids with strongly similar properties; (.) conservation between amino acids with weakly similar properties.

**S10 Table. Generation of truncated L constructs.** RdRp: RNA-dependent RNA polymerase, PRNTase: polyribonucleotidyl transferase, CD: connecting domain, MTase: methyltransferase, CTD: C-terminal domain.

